# Capturing diverse microbial sequence with comprehensive and scalable probe design

**DOI:** 10.1101/279570

**Authors:** Hayden C. Metsky, Katherine J. Siddle, Adrianne Gladden-Young, James Qu, David K. Yang, Patrick Brehio, Andrew Goldfarb, Anne Piantadosi, Shirlee Wohl, Amber Carter, Aaron E. Lin, Kayla G. Barnes, Damien C. Tully, Björn Corleis, Scott Hennigan, Giselle Barbosa-Lima, Yasmine R. Vieira, Lauren M. Paul, Amanda L. Tan, Kimberly F. Garcia, Leda A. Parham, Ikponmwonsa Odia, Philomena Eromon, Onikepe A. Folarin, Augustine Goba, Viral Hemorrhagic Fever Consortium, Etienne Simon-Lorière, Lisa Hensley, Angel Balmaseda, Eva Harris, Douglas Kwon, Todd M. Allen, Jonathan A. Runstadler, Sandra Smole, Fernando A. Bozza, Thiago M. L. Souza, Sharon Isern, Scott F. Michael, Ivette Lorenzana, Lee Gehrke, Irene Bosch, Gregory Ebel, Donald Grant, Christian Happi, Daniel J. Park, Andreas Gnirke, Pardis C. Sabeti, Christian B. Matranga

## Abstract

Metagenomic sequencing has the potential to transform microbial detection and characterization, but new tools are needed to improve its sensitivity. We developed CATCH (Compact Aggregation of Targets for Comprehensive Hybridization), a computational method to enhance nucleic acid capture for enrichment of diverse microbial taxa. CATCH designs compact probe sets that achieve full coverage of known sequence diversity and that scale well with this diversity. To illustrate applications of CATCH, we focused on capturing viral genomes. We designed, synthesized, and validated multiple probe sets, including one that targets whole genomes of the 356 viral species known to infect humans. Capture with these probe sets enriched unique viral content on average 18× and allowed us to assemble genomes that we could not otherwise recover, while accurately preserving within-sample diversity. We used this approach to recover genomes from the 2018 Lassa fever outbreak in Nigeria and to improve detection of viral infections in samples with unknown content. Together, this work demonstrates a path toward more sensitive, cost-effective metagenomic sequencing.

## Introduction

Sequencing of patient samples has revolutionized the detection and characterization of important human viral pathogens^1^ and has enabled crucial insights into their evolution and epidemiology^2–6^. Unbiased metagenomic sequencing is particularly useful for identifying and obtaining genome sequences of emerging or diverse species because it allows accurate detection of species and variants whether they are known or novel^1^. However, in practice its utility is often limited because of extremely low viral titers, e.g., as seen in the recent Zika virus outbreak^7–9^, or high levels of host material^10^. The low ratio of viral to host material results in few viral-derived sequencing reads, which can make genome assembly, if even attainable, prohibitively expensive. To fully realize the potential of metagenomic sequencing, we need new tools that improve its sensitivity while preserving its comprehensive, unbiased scope.

Previous studies have used targeted amplification^11, 12^ or enrichment via capture of viral nucleic acid using oligonucleotide probes^13–15^ to improve the sensitivity of sequencing for specific viruses. However, achieving comprehensive sequencing of viruses — similar to the use of microarrays for differential detection^16–22^ — is challenging due to the enormous diversity of viral genomes. One recent study used a probe set to target a large panel of viral species simultaneously, but did not attempt to cover strain diversity^23^. Other studies have designed probe sets to more comprehensively target viral diversity and tested their performance^24, 25^. These overcome the primary limitation of single virus enrichment methods, i.e., having to know *a priori* the taxon of interest. However, existing probe sets that target viral diversity have been designed with ad hoc approaches and they are not publicly available.

To enhance capture of diverse targets, we instead need rigorous methods, implemented in publicly available software, that can be systematically applied to create and rapidly update optimally designed probe sets. These methods ought to comprehensively cover known sequence diversity, ideally with theoretical guarantees, especially given the exceptional variability of viral genomes. Moreover, as the diversity of known taxa expands and novel species continue to be identified^26, 27^, probe sets designed by such methods must also be dynamic and scalable to keep pace with these changes. These methods should be applicable to any taxa, including all microbes. Several existing approaches to probe design for non-microbial targets^28–30^ strive to meet some of these goals but are not designed to be applied against the extensive diversity seen within and across microbial taxa.

Here, we developed and implemented CATCH (Compact Aggregation of Targets for Comprehensive Hybridization), a method that yields scalable and comprehensive probe designs from any collection of target sequences. Using CATCH, we designed several multi-virus probe sets, and then synthesized and used them to enrich viral nucleic acid in sequencing libraries from patient and environmental samples across diverse source material. We evaluated their performance and investigated any biases introduced by capture with these probe sets. Finally, to demonstrate use in clinical and biosurveillance settings, we applied this platform to recover Lassa virus genomes in low titer clinical samples from the 2018 Lassa fever outbreak in Nigeria and to identify viruses in human and mosquito samples with unknown content.

## Results

### Probe design using CATCH

To design probe sets, CATCH accepts any collection of sequences that a user seeks to target. This typically represents all known genomic diversity of one or more species. CATCH designs a set of sequences for oligonucleotide probes using a model for determining whether a probe hybridizes to a region of target sequence (Supplementary Fig. 1a; see Methods for details); the probes designed by CATCH have guarantees on capturing input diversity under this model.

CATCH searches for an optimal probe set given a desired number of oligonucleotides to output, which might be determined by factors such as cost or synthesis constraints. The input to CATCH is one or more datasets, each composed of sequences of any length, that need not be aligned to each other. In this study, each dataset consists of genomes from one species, or closely related taxa, we seek to target. CATCH incorporates various parameters that govern hybridization (Supplementary Fig. 1b), such as sequence complementarity between probe and target, and accepts different values for each dataset (Supplementary Fig. 1c). This allows, for example, more diverse datasets to be assigned less stringent conditions than others. Assume we have a function *s*(*d*, *θ*_***d***_) that gives a probe set for a single dataset *d* using hybridization parameters *θ*_***d***_, and let *S*({*θ*_***d***_}) represent the union of *s*(*d*, *θ*_***d***_) across all datasets *d* where {*θ*_***d***_} is the collection of parameters across all datasets. CATCH calculates *S*({*θ*_***d***_}), or the final probe set, by minimizing a loss function over {*θ*_***d***_} while ensuring that the number of probes in *S*({*θ*_***d***_}) falls within the specified oligonucleotide limit (Fig. 1a).

**Figure 1.**
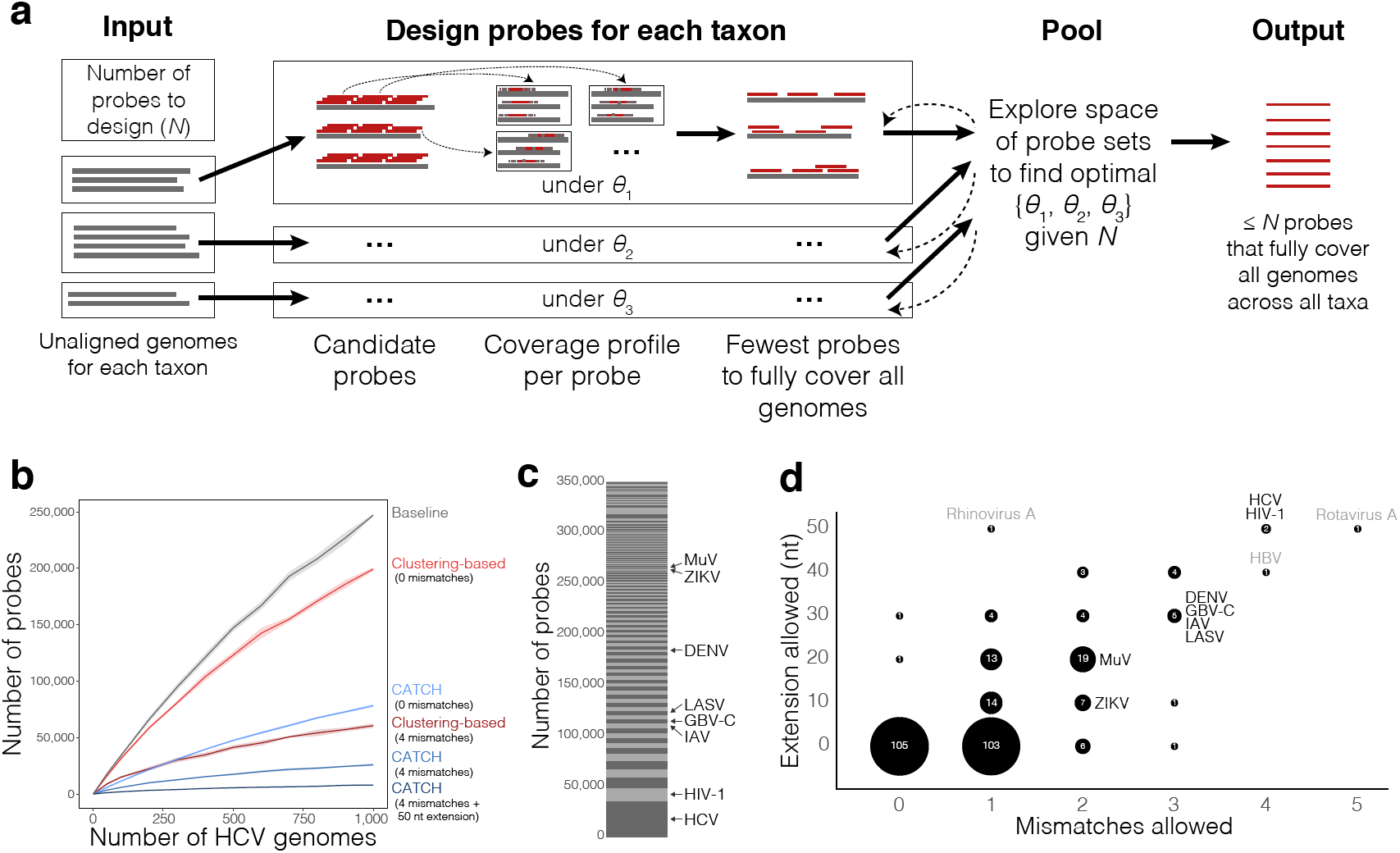
Using CATCH for probe set design. **(a)** Sketch of CATCH’s approach to probe design, shown with three datasets (typically, each is a taxon). For each dataset *d*, CATCH generates candidate probes by tiling across input genomes and, optionally, reduces the number of them using locality-sensitive hashing. Then, it determines a profile of where each candidate probe will hybridize (the genomes and regions within them) under a model with parameters *θ*_***d***_ (see Supplementary Fig. 1b for details). Using these coverage profiles, it approximates the smallest collection of probes that fully captures all input genomes (described in text as *s*(*d*, *θ*_***d***_)). Given a constraint on the total number of probes (*N*) and a loss function over the *θ*_***d***_, it searches for optimal *θ*_***d***_. **(b)** Number of probes required to fully capture increasing numbers of HCV genomes. Approaches shown are simple tiling (gray), a clustering-based approach at two levels of stringency (red), and CATCH with three choices of parameter values specifying varying levels of stringency (blue). See Methods for details regarding parameter choices. Previous approaches for targeting viral diversity use clustering in probe set design. Shaded regions around each line are 95% pointwise confidence bands calculated across randomly sampled input genomes. **(c)** Number of probes designed by CATCH for each dataset (of 296 datasets in total) among all 349,998 probes in the V**_ALL_** probe set. Species incorporated in our sample testing are labeled. **(d)** Values of the two parameters selected by CATCH for each dataset in the design of V**_ALL_**: number of mismatches to tolerate in hybridization and length of the target fragment (in nt) on each side of the hybridized region assumed to be captured along with the hybridized region (cover extension). The label and size of each bubble indicate the number of datasets that were assigned a particular combination of values. Species included in our sample testing are labeled in black, and outlier species not included in our testing are in gray. In general, more diverse viruses (e.g., HCV and HIV-1) are assigned more relaxed parameter values (here, high values) than less diverse viruses, but still require a relatively large number of probes in the design to cover known diversity (see (c)). Panels similar to (c) and (d) for the design of V**_WAFR_** are in Supplementary Fig. 3.

The key to determining the final probe set is then to find an optimal probe set *s*(*d*, *θ*_***d***_) for each input dataset. Briefly, CATCH creates “candidate” probes from the target genomes in *d* and seeks to approximate, under *θ*_***d***_, the smallest set of candidates that achieve full coverage of the target genomes. Our approach treats this problem as an instance of the well-studied *set cover problem*^31, 32^, the solution to which is *s*(*d*, *θ*_***d***_) (Fig. 1a; see Methods for details). We found that this approach scales well with increasing diversity of target genomes and produces substantially fewer probes than previously used approaches (Fig. 1b, Supplementary Fig. 2).

CATCH’s framework offers considerable flexibility in designing probes for various applications. For example, a user can customize the model of hybridization that CATCH uses to determine whether a candidate probe will hybridize to and capture a particular target sequence. Also, a user can design probe sets for capturing only a specified fraction of each target genome and, relatedly, for targeting regions of the genome that distinguish similar but distinct subtypes. CATCH also offers an option to blacklist sequences, e.g., highly abundant ribosomal RNA sequences, so that output probes are unlikely to capture them. CATCH can use locality-sensitive hashing^33, 34^, if desired, to reduce the number of candidate probes that are explored, improving runtime and memory usage on especially large numbers of input sequences. We implemented CATCH in a Python package that is publicly available at https://github.com/broadinstitute/catch.

### Probe sets to capture viral diversity

We used CATCH to design a probe set that targets all viral species reported to infect humans (V**_ALL_**), which could be used to achieve more sensitive metagenomic sequencing of viruses from human samples. V**_ALL_** encompasses 356 species (86 genera, 31 families), and we designed it using genomes available from NCBI GenBank^35, 36^ (Supplementary Table 1). We constrained the number of probes to 350,000, significantly fewer than the number used in studies with comparable goals^24, 25^, reducing the cost of synthesizing probes that target diversity across hundreds of viral species. The design output by CATCH contained 349,998 probes (Fig. 1c). This design represents comprehensive coverage of the input sequence diversity under conservative choices of parameter values, e.g., tolerating few mismatches between probe and target sequence (Fig. 1d). To compare the performance of V**_ALL_** against probe sets with lower complexity, we separately designed three focused probe sets for commonly co-circulating viral infections: measles and mumps viruses (V**_MM_**; 6,219 probes), Zika and chikungunya viruses (V**_ZC_**; 6,171 probes), and a panel of 23 species (16 genera, 12 families) circulating in West Africa (V**_WAFR_**; 44,995 probes) (Supplementary Fig. 3, Supplementary Table 1). These probe sets are publicly available (see Methods for link).

We synthesized V**_ALL_** as 75 nt biotinylated ssDNA and the focused probe sets (V**_WAFR_**, V**_MM_**, V**_ZC_**) as 100 nt biotinylated ssRNA. The ssDNA probes in V**_ALL_** are more stable and therefore more suitable for use in lower resource settings compared to ssRNA probes. We expect the ssRNA probes to be more sensitive than ssDNA probes in enriching target cDNA due to their longer length and the stronger bonds formed between RNA and DNA^37^, making the focused probe sets a useful benchmark for the performance of V**_ALL_**.

### Enrichment of viral genomes upon capture with V**_ALL_**

To evaluate enrichment efficiency of V**_ALL_**, we prepared sequencing libraries from 30 patient and environmental samples containing at least one of 8 different viruses: dengue virus (DENV), GB virus C (GBV-C), Hepatitis C virus (HCV), HIV-1, Influenza A virus (IAV), Lassa virus (LASV), mumps virus (MuV), and Zika virus (ZIKV) (see Supplementary Table 2 for details). These 8 viruses together reflect a range of typical viral titers in biological samples, including ones that have extremely low levels, such as ZIKV^38, 39^. The samples encompass a range of source materials: plasma, serum, buccal swabs, urine, avian swabs, and mosquito pools. We performed capture on these libraries and sequenced them both before and after capture. To compare enrichment of viral content across sequencing runs, we downsampled raw read data from each sample to the same number of reads (200,000) before further analysis. Downsampling to correct for differences in sequencing depth, rather than the more common use of a normalized count such as reads per million, is useful for two reasons. First, it allows us to compare our ability to assemble genomes (e.g., owing to capture) in samples that were sequenced to different depths. Second, downsampling helps to correct for differences in sequencing depth in the presence of a high frequency of PCR duplicate reads (see Methods for details), as observed in captured libraries. We removed duplicate reads during analyses so that we could measure enrichment of viral information (i.e., unique viral content) rather than measure an artifactual enrichment arising from PCR amplification.

We first assessed enrichment of viral content by examining the change in per-base read depth resulting from capture with V**_ALL_**. Overall, we observed a median increase in unique viral reads across all samples of 18 × (*Q*_1_ = 4.6, *Q*_3_ = 29.6) (Supplementary Table 3). Capture increased depth across the length of each viral genome, with no apparent preference in enrichment for regions over this length (Fig. 2a, b, Supplementary Fig. 4). Moreover, capture successfully enriched viral content in each of the 6 sample types we tested. The increase in coverage depth varied between samples, likely in part because the samples differed in their starting concentration and, as expected, we saw lower enrichment in samples with higher abundance of virus before capture (Supplementary Fig. 5).

**Figure 2.**
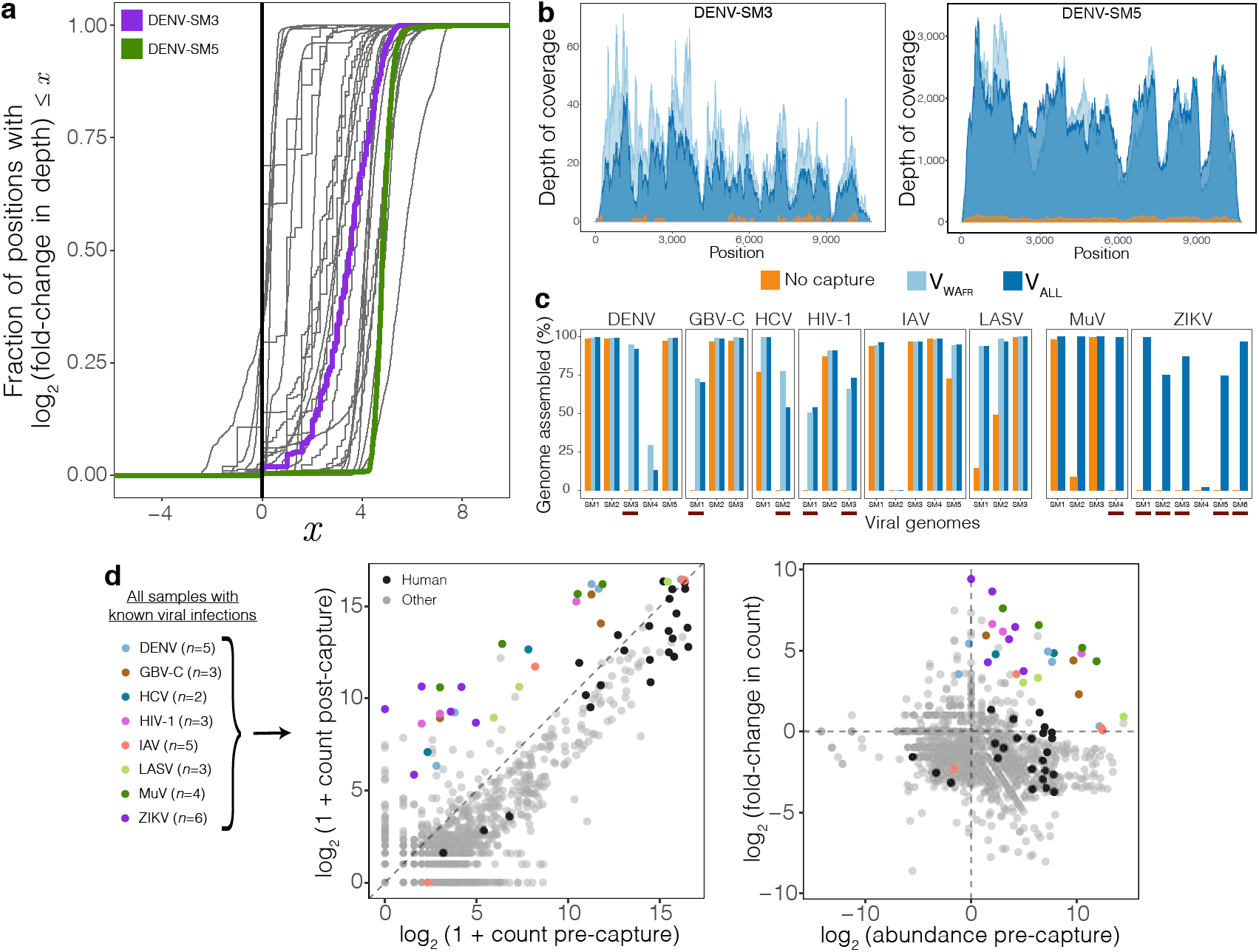
Improvement in genome coverage and assembly, and shift in metagenomic distribution after capture. **(a)** Distribution of the enrichment in read depth, across viral genomes, provided by capture with V**_ALL_** on 30 patient and environmental samples with known viral infections. Each curve represents one of the 31 viral genomes sequenced here; one sample contained two known viruses. At each position across a genome, the post-capture read depth is divided by the pre-capture depth, and the plotted curve is the empirical cumulative distribution of the log of these fold-change values. A curve that rises fully to the right of the black vertical line illustrates enrichment throughout the entirety of a genome; the more vertical a curve, the more uniform the enrichment. Read depth across viral genomes DENV-SM3 (purple) and DENV-SM5 (green) are shown in more detail in (b). **(b)** Read depth throughout a genome of DENV in two samples. DENV-SM3 (left) has few informative reads before capture and does not produce a genome assembly, but does following capture. DENV-SM5 (right) does yield a genome assembly before capture, and depth increases following capture. **(c)** Percent of the viral genomes unambiguously assembled in the 30 samples, which had 8 known viral infections across them. Shown before capture (orange), after capture with V**_WAFR_** (light blue), and after capture with V**_ALL_** (dark blue). Red bars below samples indicate ones in which we could not assemble any contig before capture but, following capture, were able to assemble at least a partial genome (> 50%). **(d)** Left: Number of reads detected for each species across the 30 samples with known viral infections, before and after capture with V**_ALL_**. Reads in each sample were downsampled to 200,000 reads. Each point represents one species detected in one sample. For each sample, the virus previously detected in the sample by another assay is colored. *Homo sapiens* matches in samples from humans are shown in black. Right: Abundance of each detected species before capture and fold-change upon capture with V**_ALL_** for these samples. Abundance was calculated by dividing pre-capture read counts for each species by counts in pooled water controls. Coloring of human and viral species are as in the left panel.

Next we analyzed how capture improved our ability to assemble viral genomes. For samples that had partial genome assemblies (< 90%) before capture, we found that application of V**_ALL_** allowed us to assemble a greater fraction of the genome in all cases (Fig. 2c). Importantly, of the 14 samples from which we were unable to assemble any contig before capture, 11 assembled at least partial genomes (> 50%) using V**_ALL_**, of which 4 were complete genomes (> 90%). Many of the viruses we tested, such as HCV and HIV-1, are known to have high within-species diversity yet the enrichment of their unique content was consistent with that of less diverse species (Supplementary Table 3).

We also explored the impact of capture on the complete metagenomic diversity within each sample. Metagenomic sequencing generates reads from the host genome as well as background contaminants^40^, and capture ought to reduce the abundance of these taxa. Following capture with V**_ALL_**, the fraction of sequence classified as human decreased in patient samples while viral species with a wide range of pre-capture abundances were strongly enriched (Fig. 2d). Moreover, we observed a reduction in the overall number of species detected after capture (Supplementary Fig. 6a), suggesting that capture indeed reduces non-targeted taxa. Lastly, analysis of this metagenomic data identified a number of other enriched viral species present in these samples (Supplementary Table 4). For example, one HIV-1 sample showed strong evidence of HCV co-infection, an observation consistent with clinical PCR testing.

In addition to measuring enrichment on patient and environmental samples, we sought to evaluate the sensitivity of V**_ALL_** on samples with known quantities of viral and background material. To do so, we performed capture with V**_ALL_** on serial dilutions of Ebola virus (EBOV) — ranging from 10^6^ copies down to single copy — in known background amounts of human RNA. At a depth of 200,000 reads, use of V**_ALL_** allowed us to reliably detect viral content (i.e., observe viral reads in two technical replicates) down to 100 copies in 30 ng of background and 1,000 copies in 300 ng (Fig. 3a, Supplementary Table 5), each at least an order of magnitude fewer than without capture, and similarly lowered the input at which we could assemble genomes (Supplementary Fig. 7a). Although we chose a single sequencing depth so that we could compare pre- and post-capture results, higher sequencing depths provide more viral material and thus more sensitivity in detection (Supplementary Fig. 7b, c).

**Figure 3.**
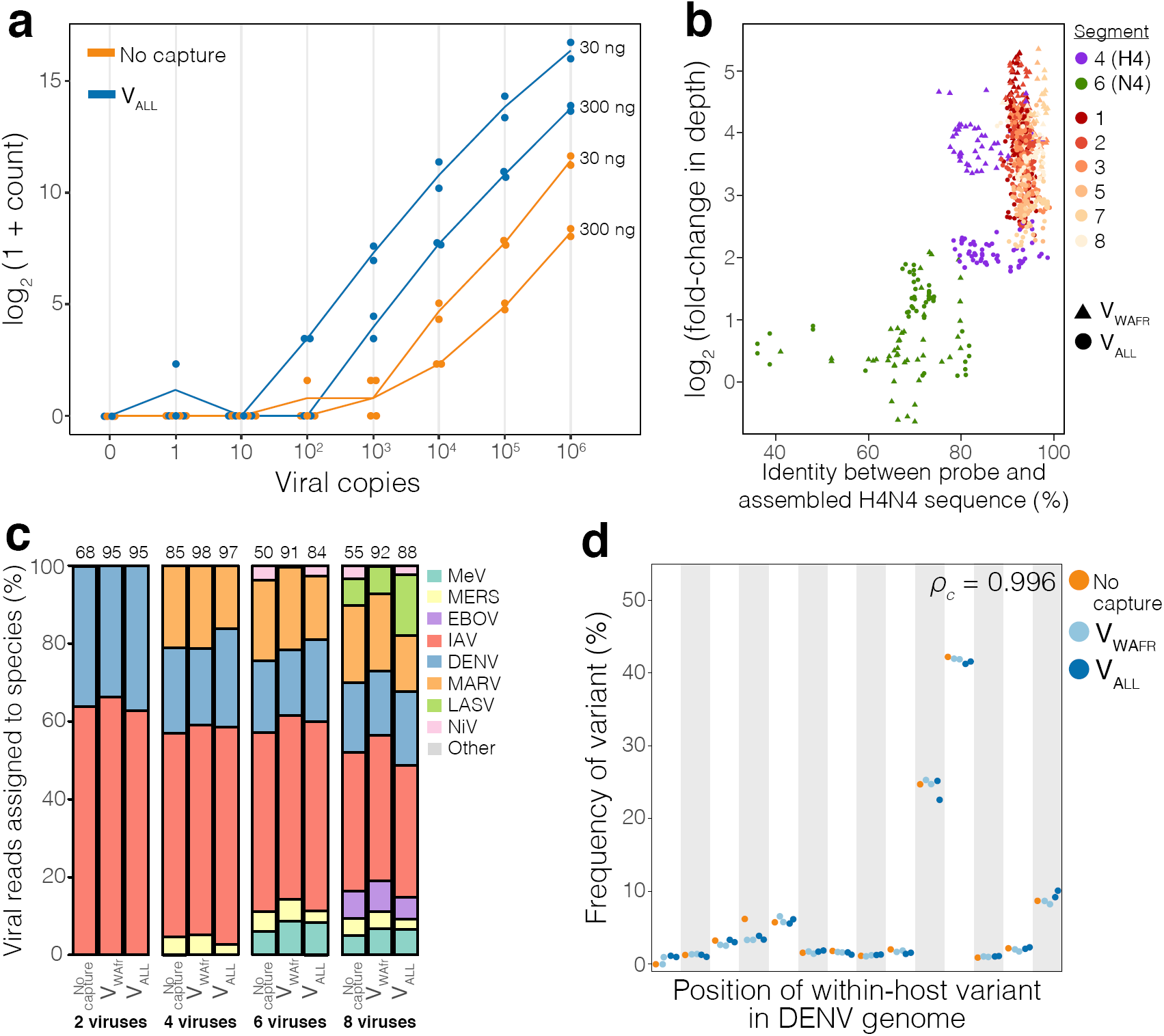
Characterizing improvement in detection and detection of within-sample diversity. **(a)** Amount of viral material sequenced in a dilution series of viral input in two amounts of human RNA background. There are *n* = 2 technical replicates for each choice of input copies, background amount, and use of capture (*n* = 1 replicate for the negative control with 0 copies). Each dot indicates number of reads, among 200,000 in total, sequenced from a replicate; line is through the mean of the replicates. Label to the right of each line indicates amount of background material. **(b)** Relation between probe-target identity and enrichment in read depth, as seen after capture with V**_ALL_** and with V**_WAFR_** on an Influenza A virus sample of subtype H4N4 (IAV-SM5). Each point represents a window in the IAV genome. Identity between the probe and assembled H4N4 sequence is a measure of identity between the sequence in that window and the top 25% of probe sequences that map to it (see Methods for details). Fold-change in depth is averaged over the window. No sequences of segment 6 (N) of the N4 subtypes were included in the design of V**_ALL_** or V**_WAFR_**. **(c)** Effect of capture on estimated frequency of within-sample co-infections. RNA of 2, 4, 6, and 8 viral species were spiked into extracted RNA from healthy human plasma and then captured with V**_ALL_** and V**_WAFR_**. Values on top are the percent of all sequenced reads that are viral. We did not detect Nipah virus (NiV) using the V**_WAFR_** probe set because this virus was not present in that design. **(d)** Effect of capture on estimated frequency of within-host variants, shown in positions across three dengue virus samples: DENV-SM1, DENV-SM2, and DENV-SM5. Capture with V**_ALL_** and V**_WAFR_** was each performed on *n* = 2 replicates of the same library. *ρ*_***c***_ indicates concordance correlation coefficient between preand post-capture frequencies.

### Comparison of V**_ALL_** to focused probe sets

To test whether the performance of the highly complex 356-virus V**_ALL_** probe set matches that of focused ssRNA probe sets, we first compared it to the 23-virus V**_WAFR_** probe set. We evaluated the 6 viral species we tested from the patient and environmental samples that were present in both the V**_ALL_** and V**_WAFR_** probe sets, and we found that performance was concordant between them: V**_WAFR_** provides almost the same number of unique viral reads as V**_ALL_** (1.01× as many; *Q*_1_ = 0.93, *Q*_3_ = 1.34) (Supplementary Table 3). The percentage of each genome that we could unambiguously assemble was also similar between the probe sets (Fig. 2c), as was the read depth (Supplementary Fig. 4, Supplementary Fig. 8a, b). Following capture with V**_WAFR_**, human material and the overall number of detected species both decreased, as with V**_ALL_**, although these changes were more pronounced with V**_WAFR_** (Supplementary Fig. 6a, b, Supplementary Table 4).

We next compared the V**_ALL_** probe set to the two 2-virus probe sets V**_MM_** and V**_ZC_**. We found that enrichment for MuV and ZIKV samples was slightly higher using the 2-virus probe sets than with V**_ALL_** (2.26 × more unique viral reads; *Q*_1_ = 1.69, *Q*_3_ = 3.36) (Supplementary Table 3, Supplementary Fig. 4, Supplementary Fig. 8c, d). The additional gain of these probe sets might be useful in some applications, but was considerably less than the 18× increase provided by V**_ALL_** against a pre-capture sample. Overall, our results suggest that neither the complexity of the V**_ALL_** probe set nor its use of shorter ssDNA probes prevent it from efficiently enriching viral content.

### Enrichment of targets with divergence from design

We then evaluated how well our V**_ALL_** and V**_WAFR_** probe sets capture sequence that is divergent from sequences used in their design. To do this, we tested whether the probe sets, whose designs included human IAV, successfully enrich the genome of the non-human, avian subtype H4N4 (IAV-SM5). H4N4 was not included in the designs, making it a useful test case for this relationship. Moreover, the IAV genome has 8 RNA segments that differ considerably in their genetic diversity; segment 4 (hemagglutinin; H) and segment 6 (neuraminidase; N), which are used to define the subtypes, exhibit the most diversity.

The segments of the H4N4 genome display different levels of enrichment following capture (Supplementary Fig. 9). To investigate whether these differences are related to sequence divergence from the probes, we compared the identity between probes and sequence in the H4N4 genome to the observed enrichment of that sequence (Fig. 3b). We saw the least enrichment in segment 6 (N), which had the least identity between probe sequence and the H4N4 sequence, as we did not include any sequences of the N4 subtypes in the probe designs. Interestingly, V**_ALL_** did show limited positive enrichment of segment 6, as well as of segment 4 (H); these enrichments were lower than those of the less divergent segments. But this was not the case for segment 4 when using V**_WAFR_**, suggesting a greater target affinity of V**_WAFR_** capture when there is some degree of divergence between probes and target sequence (Fig. 3b), potentially due to this probe set’s longer, ssRNA probes. For both probe sets, we observed no clear inter-segment differences in enrichment across the remaining segments, whose sequences have high identity with probe sequences (Fig. 3b, Supplementary Fig. 9). These results show that the probe sets can capture sequence that differs markedly from what they were designed to target, but nonetheless that sequence similarity with probes influences enrichment efficiency.

### Quantifying within-sample diversity after capture: co-infections and within-host nucleotide variants

Given that many viruses co-circulate within geographic regions, we assessed whether capture accurately preserves within-sample viral species complexity. We first evaluated capture on mock co-infections containing 2, 4, 6, or 8 viruses. Using both V**_ALL_** and V**_WAFR_**, we observed an increase in overall viral content while preserving relative frequencies of each virus present in the sample (Fig. 3c, Supplementary Table 4).

Because viruses often have extensive within-host viral nucleotide variation that can inform studies of transmission and within-host virus evolution^41–44^, we examined the impact of capture on estimating within-host variant frequencies. We used three DENV samples that yielded high read depth (Supplementary Table 3). Using both V**_ALL_** and V**_WAFR_**, we found that frequencies of all within-host variants were consistent with pre-capture levels (Fig. 3d, Supplementary Table 6; concordance correlation coefficient is 0.996 for V**_ALL_** and 0.997 for V**_WAFR_**). These estimates were consistent for both low and high frequency variants. Since capture preserves frequencies so well, it should enable measurement of within-host diversity that is both sensitive and cost-effective.

### Rescuing Lassa virus genomes in patient samples from Nigeria

To demonstrate the application of V**_ALL_** in the case of an outbreak, we applied it to samples of clinically confirmed (by qRT-PCR) Lassa fever cases from Nigeria. In 2018, Nigeria experienced a sharp increase in cases of Lassa fever, a severe hemorrhagic disease caused by LASV^45^, leading the World Health Organization and the Nigeria Centre for Disease Control to declare it an outbreak^46, 47^. Previous genome sequencing of LASV has revealed its extensive genetic diversity, with distinct lineages circulating in different parts of the endemic region^3, 48^, and ongoing sequencing can enable rapid identification of changes in this genetic landscape.

We selected 23 samples, spanning 5 states in Nigeria, that yielded either no portion of a LASV genome or only partial genomes with unbiased metagenomic sequencing even at high sequencing depth (> 4.5 million reads)^49^, and performed capture on these using V**_ALL_**. At equivalent pre- and post-capture sequencing depth (200,000 reads), use of V**_ALL_** improved our ability to detect and assemble LASV. Capture considerably increased the amount of unique LASV material detected in all 23 samples (in 4 samples, by more than 100 ×), and in 7 samples it enabled detection when there were no LASV reads pre-capture (Supplementary Fig. 10a, Supplementary Table 7). This in turn improved genome assembly. Whereas precapture we could not assemble any portion of a genome in 22 samples (in the remaining one, 2% of a genome) at this depth, following use of V**_ALL_** we could assemble a partial genome in 22 of the 23 (Fig. 4a, Supplementary Fig. 10b); most were small portions of a genome, although in 7 we assembled > 50% of a genome. Assembly results with V**_ALL_** are comparable without downsampling (Supplementary Fig. 10c), likely because we saturate unique content with V**_ALL_** even at low sequencing depths (Supplementary Fig. 7b, c). These results illustrate how V**_ALL_** can be used to improve viral detection and genome assembly in an outbreak, especially at the low sequencing depths that may be desired or required in these settings.

**Figure 4.**
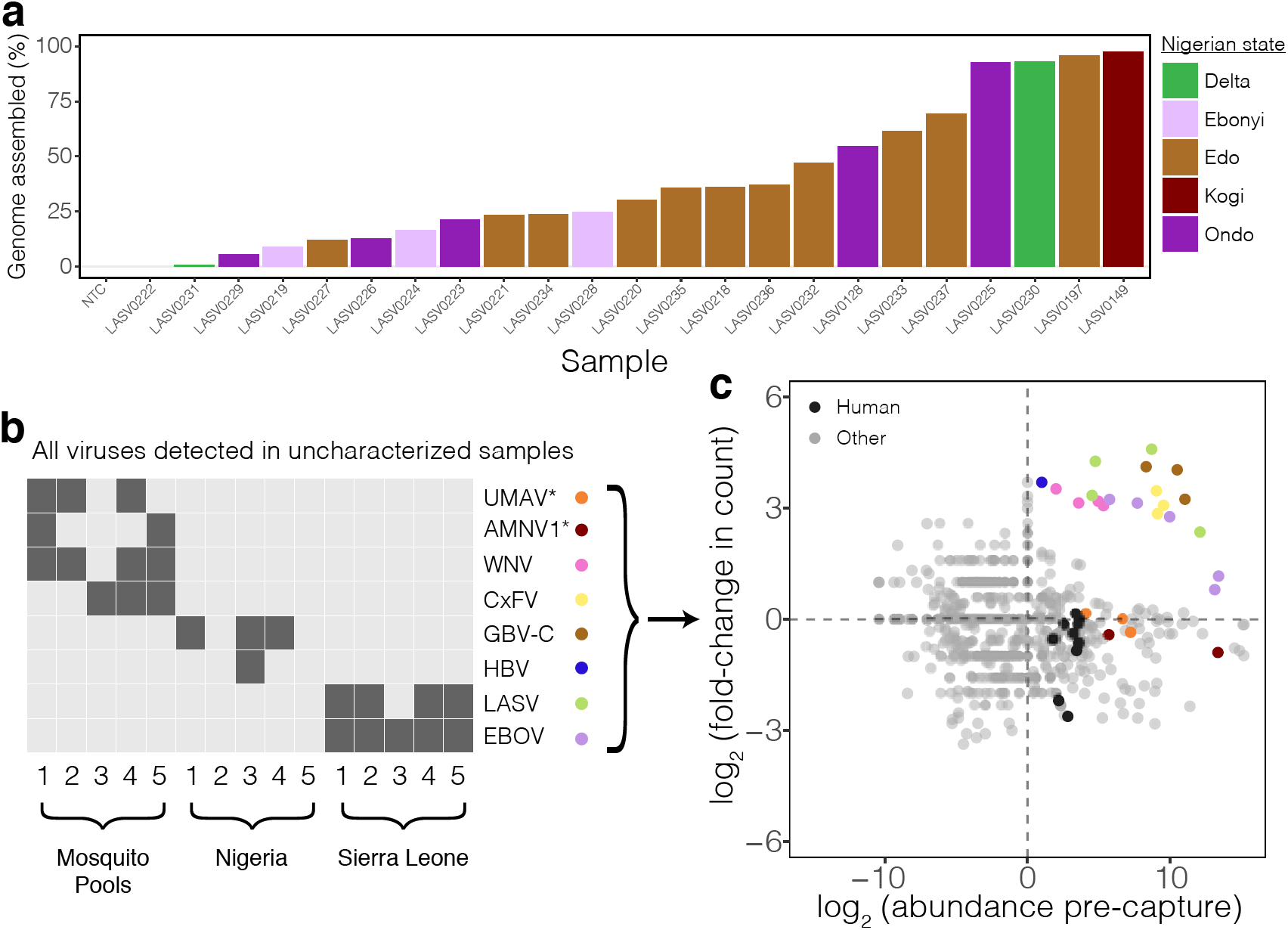
Genomic applications using capture: sequencing from the 2018 Lassa fever outbreak and of infections in uncharacterized samples. **(a)** Percent of LASV genome assembled, after use of V**_ALL_**, among 23 samples from the 2018 Lassa fever outbreak. Reads were downsampled to 200,000 reads before assembly. Bars are ordered by amount assembled and colored by the state in Nigeria that the sample is from. **(b)** Viral species present in uncharacterized mosquito pools and pooled human plasma samples from Nigeria and Sierra Leone after capture with V**_ALL_**. Asterisks on species indicate ones that are not targeted by V**_ALL_**. Detected viruses include Umatilla virus (UMAV), Alphamesonivirus 1 (AMNV1), West Nile virus (WNV), Culex flavivirus (CxFV), GBV-C, Hepatitis B virus (HBV), LASV, and EBOV. **(c)** Abundance of all detected species before capture and fold-change upon capture with V**_ALL_** in the uncharacterized sample pools. Abundance was calculated as described in Fig. 2d. Viral species present in each sample (see (b)) are colored, and *Homo sapiens* matches in the human plasma samples are shown in black.

### Identifying viruses in uncharacterized samples using capture

We next applied our V**_ALL_** probe set to pools of human plasma and mosquito samples with uncharacterized infections. We tested 5 pools of human plasma from a total of 25 individuals with suspected LASV or EBOV infections from Sierra Leone, as well as 5 pools of human plasma from a total of 25 individuals with acute fevers of unknown cause from Nigeria and 5 pools of *Culex tarsalis* and *Culex pipiens* mosquitoes from the United States (see Methods for details). Using V**_ALL_** we detected 8 viral species, each present in one or more pools: 2 species in the pools from Sierra Leone, 2 species in the pools from Nigeria, and 4 species in the mosquito pools (Fig. 4b, Supplementary Fig. 6c). We found consistent results with V**_WAFR_** for the species that were included in its design (Supplementary Fig. 6d, Supplementary Table 4). To confirm the presence of these viruses we assembled their genomes and evaluated read depth (Supplementary Fig. 11, Supplementary Table 8). We also sequenced pre-capture samples and saw significant enrichment by capture (Fig. 4c, Supplementary Fig. 6c, d). Quantifying abundance and enrichment together provides a valuable way to discriminate viral species from other taxa (Fig. 4c), thereby helping to uncover which pathogens are present in samples with unknown infections.

Looking more closely at the identified viral species, all pools from Sierra Leone contained LASV or EBOV, as expected (Fig. 4b). The 5 plasma pools from Nigeria showed little evidence for pathogenic viral infections; however, one pool did contain Hepatitis B virus. Additionally, 3 pools contained GBV-C, consistent with expected frequencies for this region^26, 50^. In mosquitoes, 4 pools contained West Nile virus (WNV), a common mosquito-borne infection, consistent with PCR testing. In addition, 3 pools contained Culex flavivirus, which has been shown to co-circulate with WNV and co-infect *Culex* mosquitoes in the United States^51^. These findings demonstrate the utility of capture to improve virus identification without *a priori* knowledge of sample content.

## Discussion

In recent years metagenomic sequencing has been widely used to investigate viral infections and outbreaks, but is often limited in practice due to low sensitivity. Capture using oligonucleotide probes to enrich viral content is one approach that can address this limitation^13–15, 23–25^. Here we describe CATCH, a method that condenses highly diverse target sequence data into a small number of oligonucleotides, enabling more efficient and sensitive sequencing that is only biased by the extent of known diversity. We show that capture with probe sets designed by CATCH improved viral genome detection and recovery, across a range of sample source materials, while accurately preserving sample complexity of the targets. Probe sets we present here have also helped us to assemble genomes of low titer viruses in other patient samples: V**_ZC_** for suspected ZIKV cases^7^ and V**_ALL_** for improving rapid detection of Powassan virus in a clinical case^52^.

The probe sets we have designed with CATCH, and more broadly capture with comprehensive probe designs, improve the accessibility of metagenomic sequencing in resource-limited settings through smaller capacity platforms. For example, in West Africa we are using the V**_ALL_** probe set to characterize LASV and other viruses in patients with undiagnosed fevers by sequencing on a MiSeq (Illumina). This could also be applied on other small machines such as the iSeq (Illumina) or MinION (Oxford Nanopore)^53^. Further, the increase in viral content enables more samples to be pooled and sequenced on a single run, increasing sample throughput and decreasing per-sample cost relative to unbiased sequencing (Supplementary Table 9). Lastly, researchers can use CATCH to quickly design focused probe sets, providing flexibility when it is not necessary to target an exhaustive list of viruses, such as in outbreak response or for targeting pathogens associated with specific clinical syndromes.

Despite the potential of capture, there are challenges and practical considerations that are present with the use of any probe set. Notably, as capture requires additional cycles of amplification, computational analyses should properly account for duplicate reads due to amplification; the inclusion of unique molecular identifiers^54, 55^ could improve determination of unique fragments. Also, quantifying the sensitivity and specificity of capture with comprehensive probe sets is challenging — as it is for metagenomic sequencing more broadly — because doing so would necessitate obtaining viral genomes for the hundreds of targeted species, and false positives are likely to be due to components of sequencing and classification that are unrelated to capture (e.g., contamination in sample processing or read misclassifications). For sequencing some ultra low input samples, targeted amplicon approaches may be faster and more sensitive^38^, but genome size, sequence heterogeneity, and the need for prior knowledge of the target species can limit the feasibility and sensitivity of these approaches^1, 56, 57^. Similarly, for molecular diagnostics of particular pathogens, many commonly used assays such as qRT-PCR and rapid antigen tests are likely to be faster and less expensive than metagenomic sequencing. Capture does increase the preparation cost and time per-sample compared to unbiased metagenomic sequencing, but this is offset by reduced sequencing costs through increased sample pooling and/or lower-depth sequencing^1^ (Supplementary Table 9).

While our conclusions here are based on whole genome capture of viruses, CATCH is a versatile approach that can be applied to capture of non-viral microbial genomes and to the design of oligonucleotide sequences for uses other than whole genome enrichment. Because CATCH scales well with our growing knowledge of genomic diversity^26, 27^, it is particularly wellsuited for designing against any class of input from microbes that have a high degree of diversity. Capture-based approaches have successfully been used to enrich whole genomes of eukaryotic parasites such as Plasmodium^58^ and Babesia^59^, as well as bacteria^60^. Many bacteria, like viruses, have high variation even within species^61^, and CATCH can enable efficient and sensitive enrichment of these bacterial genomes or even of combinations of viral and bacterial targets. Beyond microbes, CATCH can benefit studies in other areas that use capture-based approaches, such as the detection of previously characterized fetal and tumor DNA from cell-free material^62, 63^, in which known targets of interest may represent a small fraction of all material and for which it may be useful to rapidly design new probe sets for enrichment as novel targets are discovered. Moreover, CATCH can identify conserved regions or regions suitable for differential identification, which can help in the design of PCR primers and CRISPR-Cas13 crRNA guides for nucleic acid diagnostics.

CATCH is, to our knowledge, the first approach to systematically design probe sets for whole genome capture of highly diverse target sequences that span many species. Our results show that it offers an important extension to the field’s toolkit for effective viral detection and surveillance with enrichment and other targeted approaches. We anticipate that CATCH, together with these approaches, will help provide a more complete understanding of genetic diversity across the microbial world.

## Acknowledgements

We thank S. Ye, C. Myhrvold, S. Weingarten-Gabbay, C. Freije, S. Schaffner, and other members of the Sabeti Laboratory for useful discussions and feedback on the manuscript; B. Chak for assistance with ethical approvals and compliance; and Boca Biolistics, the Florida Department of Health, Miami-Dade County Mosquito Control, Research Blood Components, the Ragon Institute Cellular Immunology Database, and Brigham and Women’s Hospital’s Crimson Core for support with samples. This project has been funded in whole or in part with Federal funds from the National Institute of Allergy and Infectious Diseases, National Institutes of Health, Department of Health and Human Services, under Grant Number U19AI110818 to the Broad Institute. This project was also funded in part by NIH NIAID contract HHSN272200900049C, a Broad*next*10 gift from the Broad Institute, the Henry M. Jackson Foundation award W81XWH-11-2-0174, and the Bill & Melinda Gates Foundation. IAV samples were funded by NIH NIAID contract HHSN272201400008C to J.A.R. K.J.S. is supported by a fellowship from the Human Frontiers in Science Program (LT000553/2016). S.I. and S.F.M. are supported by NIH NIAID R01AI099210. C.H is supported by NIH NHGRI U01HG007480 and U54HG007480, and World Bank project ACE019. H.C.M., D.J.P., A.Gn., P.C.S. and C.B.M. are co-inventors on a patent application filed by the Broad Institute related to work in this manuscript (WO/2017/040316).

## Author contributions

H.C.M., D.J.P., A.Gn., P.C.S., and C.B.M. initiated the study of improved design and application of comprehensive probe sets.

H.C.M. conceived of CATCH and implemented it with advice from D.J.P., A.Gn., and C.B.M.

K.J.S. and C.B.M. conceived of experimental design for evaluating probe sets.

C.B.M., J.Q., A.G.-Y., and K.J.S. developed enrichment protocols with help from A.Gol.

K.J.S, A.G.-Y., J.Q., and P.B. prepared samples, performed enrichment, and sequenced samples.

A.P., S.W., A.C., A.L., and K.G.B. helped with sample preparation and enrichment.

D.C.T., B.C., S.H., G.B.-L., Y.R.V., L.M.P., A.L.T., K.F.G., L.A.P., A.B., E.H., D.K., T.M.A., J.A.R., S.S., T.M.L.S., S.I., S.F.M., I.L., L.G., and I.B. collected and shared samples with known viral content.

E.S.-L. and L.H. shared viral seed stocks.

G.E. shared uncharacterized mosquito pools.

I.O., P.E., O.A.F., A.Gob., D.G., and C.H. collected human plasma samples from Nigeria and Sierra Leone.

H.C.M. and K.J.S. formulated and performed data analyses with help from D.K.Y. H.C.M., K.J.S., and C.B.M. wrote the manuscript with input from other authors.

## Methods

### Probe design using CATCH

#### Designing a probe set given a single choice of parameters

We first describe how CATCH determines a probe set that covers input sequences under some selection of parameters. That is, the input is a collection of (unaligned) sequences *d* and parameters *θ*_***d***_ describing hybridization, and the goal is to compute a set of probes *s*(*d*, *θ*_***d***_). For example, *d* commonly encompasses the strain diversity of one or more species and *θ*_***d***_ includes the number of mismatches that we ought to tolerate when determining whether a probe hybridizes to a sequence.

CATCH produces a set of “candidate” probes from the input sequences in *d* by stepping along them according to a specified stride (Fig. 1a). Optionally, CATCH uses locality-sensitive hashing^33, 34^ (LSH) to reduce the number of candidate probes, which is particularly useful when the input is a large number of highly similar sequences. CATCH supports two LSH families: one under Hamming distance^33^ and another using the MinHash technique^34, 64^, which has been used in metagenomic applications^65, 66^. It detects near-duplicate candidate probes by performing approximate near neighbor search^34^ using a specified family and distance threshold. CATCH constructs hash tables containing the candidate probes and then queries each (in descending order of multiplicity) to find and collapse near-duplicates. Because LSH reduces the space of candidate probes, it may remove candidate probes that would otherwise be selected in steps described below, thereby increasing the size of the output probe set. Use of LSH to reduce the number of candidate probes is optional in our implementation of CATCH; we did not use it to produce the probe sets in this work. The approach of detecting near-duplicates among probes (and subsequently mapping them onto sequences, described below) bears some similarity to the use of *P-clouds* for clustering related oligonucleotides in order to identify diverse repetitive regions in the human genome^67, 68^.

CATCH then maps each candidate probe *p* back to the target sequences with a seed-andextend-like approach, in the process deciding whether *p* maps to a range *r* in a target sequence according to a function *f*_map_(*p*, *r*, *θ*_***d***_). *f*_map_ effectively specifies whether *p* will capture the subsequence at *r*. Further, CATCH assumes that because *p* captures an entire fragment and not just the subsequence to which it binds, *p* “covers” both *r* and some number of bases (given in *θ*_***d***_) on each side of *r*; we term this a “cover extension”. This yields a collection of bases in the target sequences that are covered by each *p*, namely:

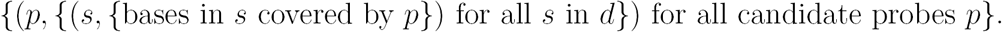

Next, CATCH seeks to find the smallest set of candidate probes that achieves full coverage of all sequences in *d*. The problem is NP-hard. To determine *s*(*d*, *θ*_***d***_), an approximation of the smallest such set of candidates probes, CATCH treats the problem as an instance of the set cover problem. Similar approaches have been used in related problems in uncovering patterns in DNA sequence. Notably, these include PCR primer selection^69–71^, string barcoding of pathogens^72, 73^, and other applications in microbial microarrays^74–76^, although these are not aimed at whole genome enrichment for sequencing many taxa.

CATCH computes *s*(*d*, *θ*_***d***_) using the canonical greedy solution to the set cover problem^31^,32, which likely provides close to the best achievable approximation^77^. In this approximation-preserving reduction, each candidate probe *p* is treated as a set whose elements represent the bases in the target sequences covered by *p*. The universe of elements is then all the bases across all the target sequences — i.e., what it seeks to cover. To implement the algorithm efficiently, CATCH operates on sets of intervals rather than base positions and applies other techniques to improve performance for this particular problem.

### Extensions to probe design

This framework for designing probes offers considerable flexibility. For example, it reduces the design to a problem of determining probe-target hybridization. The function *f*_map_, which determines whether a probe hybridizes to a range in a target sequence (and, if it does, precisely the range), can be customized by a user in CATCH’s source code or can be provided in a command-line argument to be dynamically loaded. For example, although by default CATCH does not use a thermodynamic model of hybridization, a user could choose to incorporate a calculation of free energy to evaluate the likelihood of hybridization. Here, when computing *s*(*d*, *θ*_***d***_), CATCH’s default *f*_map_ is based on three parameters in *θ*_***d***_: a number *m* of mismatches to tolerate, a length *lcf* of a longest common substring, and a length *i* of an island of an exact match. *f*_map_ computes the longest common substring with at most *m* mismatches between the probe sequence and target subsequence, and returns that the probe covers the target range if and only if the length of this is at least *lcf*. Optionally (if *i* > 0), *f*_map_ additionally requires that the probe and target subsequence share an exact (0-mismatch) match of length at least *i* to return that the probe covers the range. (See Supplementary Fig. 1b for a visual representation and “Exploring the parameter space across taxa” for example values.)

There are many problems related to probe design that map well to generalizations of the set cover problem. Relevant generalizations are the weighted and partial cover problems^31, 78, 79^. Using the weighted cover problem, CATCH allows a user to perform differential identification of taxa and also to blacklist sequences from the probe design. For these purposes, we introduce the concept of a “rank” to our implementation of the set cover solution. A rank of a set is analogous to a weight and makes it straightforward to assign levels of penalties on sets. For two sets *S* and *T*, if rank(*S*) < rank(*T*) then *S* is always considered before *T* — i.e., if coverage is needed and *S* provides that coverage, then the greedy algorithm always chooses *S* before *T* even if *T* provides more. These can be emulated entirely using weights (i.e., costs), by assigning sufficiently high weights to each set. To perform differential identification, CATCH accepts groupings of sequences as input (for example, each grouping might encompass the available genomes of a species). Then, CATCH finds the number of groupings that each candidate probe *p* “hits”. (*p* hits a grouping if it covers a part of at least one sequence in that grouping.) A probe that hits only one grouping is suitable for differential identification, whereas ones that hit more are poor choices. Thus, CATCH assigns a rank to each *p* equal to the number of groupings hit by *p*. CATCH can also accept a collection of sequences to blacklist from the probe design. It determines the number of nucleotides in blacklisted sequence that each *p* covers and assigns to *p* a rank equal to this value; therefore, candidate probes that cover blacklisted sequence are highly penalized in the design. (When a user opts to perform differential identification while also blacklisting sequences, the ranks are assigned such that a candidate probe that covers a part of a blacklisted sequence always receives a higher rank than one that does not.) For the purposes of determining whether *p* hits an identification grouping or blacklisted sequence, CATCH accepts three additional parameters, holding more tolerant values for *m*, *lcf*, and *i* as defined above, that *f*_map_ uses to evaluate probe-target hybridization. We note as well that weights can have other applications in probe design, e.g., if there is a reason to prefer some candidate probes over others due to base composition. Finally, CATCH solves an instance of the weighted cover problem by assigning the rank of each set to be the rank of the candidate probe it represents.

Based on the partial cover problem, CATCH offers the ability to design probes such that they only cover a portion of each target sequence. The user specifies this portion as either a fraction of the length of each sequence or as a fixed number of nucleotides. Reducing the problem directly to an instance of the set cover problem with a single universe would not allow partially covering *each* target sequence. Thus, we introduce “multiple universes” to the instance, in which each universe corresponds to a target sequence and consists of all the bases in that sequence. Each set (representing candidate probes) specifies which elements in which universes it covers. The greedy algorithm continues selecting among the candidate probes until it obtains the desired partial coverage of each universe (target sequence). We don’t make claims about the approximation factor this achieves. As one application, note that when performing differential identification the required partial coverage should be set to be relatively low.

If desired, CATCH adds adapters to probe sequences in *s*(*d*, *θ*_***d***_) for PCR amplification. Because probe sequences may overlap, it is possible that, during PCR, they could chain together to form concatemers. Thus, we would like to use *k* unique adapters and divide the probes in *s*(*d*, *θ*_***d***_) into *k* groups such that the probes in each group are unlikely to chain together; then, we can perform PCR separately on each group. CATCH uses a heuristic to solve this problem for *k* = 2, i.e., two adapters *A* and *B*. Consider one target sequence *t*. It maps each of the probes in *s*(*d*, *θ*_***d***_) to *t* using *f*_map_, as described above. It treats the ranges that each probe covers as an “interval,” and finds the largest set of non-overlapping intervals (probes) *T*_no_ by solving an instance of the interval scheduling problem. Then, we could assign adapter *A* to each probe in *T*_no_, and adapter *B* to each of the others. CATCH performs this for each target sequence *t*, and each *t* “votes” once (either *A* or *B*) for each probe. We seek to maximize the sum, across all probes, of the majority vote for the probe (to ensure a clear decision on the adapter for each probe). Let 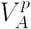 be the number of *A* votes for a probe, and likewise for 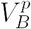. Then, we wish to maximize the quantity

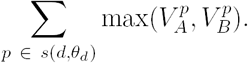

Since the distinction between *A* and *B* is arbitrary, at each *t* CATCH chooses whether to assign *A* or *B* votes to the probes in *T*_no_ depending on which assignment yields a higher sum. This process yields the maximum sum, and CATCH then assigns adapter *A* or *B* to each probe based on which has more votes.

### Designing across many taxa

Consider a large set of input sequences that encompass a diverse set of taxa (e.g., hundreds of viral species). We could run CATCH, as described above, on a single choice of parameters *θ*_***d***_ such that the number of probes in *s*(*d*, *θ*_***d***_) is feasible for synthesis. However, this can lead to a poor representation of taxa in the diverse probe set; it can become dominated by probes covering taxa that have more genetic diversity (e.g., HIV-1). Furthermore, it can force probes to be designed with relaxed assumptions about hybridization across all taxa. To alleviate these issues, we allow different choices of parameters governing hybridization for different subsets of input sequences, so that some can have probes designed with more relaxed assumptions than others.

We represent a set of taxa and its target sequences with a dataset *d*, with its own set of parameters *θ*_***d***_. Let {*θ*_***d***_} be the collection of *θ*_***d***_ across all *d*. We wish to find *S*({*θ*_***d***_}), the union of *s*(*d*, *θ*_***d***_) across all datasets *d*. CATCH finds this by solving a constrained nonlinear optimization problem:

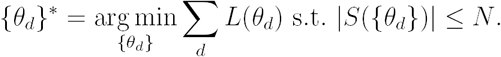

The constraint *N* on the number of probes in the union is specified by the user; this is the number of probes to synthesize, and might be determined based on synthesis cost and/or array size. CATCH solves this using the barrier method with a logarithmic barrier function. By default, we use the following loss function for each *d*:

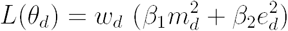

where *m*_***d***_ gives a number of mismatches to tolerate in hybridization and *e*_***d***_ gives a cover extension, as defined above. *w*_***d***_ allows a relative weighting of datasets, e.g., if one should have more stringent assumptions about hybridization and thus more probes. *β*_1_, *β*_2_, and the set of {*w*_***d***_} s can be specified by the user. A user can also choose to generalize the search to a different set of parameters:

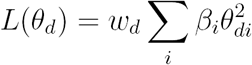

where *θ*_***di***_ is the value of the *i*th parameter for *d* and *β*_***i***_ is a specified coefficient for that parameter.

In practice, we have used the default loss function above, with *w*_***d***_ = 1 for all *d*, *β*_1_ = 1, and 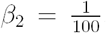. We calculate *s*(*d*, *θ*_***d***_) for each *d* over a grid of values of *θ*_***d***_ before solving for {*θ*_***d***_}*. CATCH interpolates |*s*(*d*, *θ*_***d***_)| for non-computed values of *θ*_***d***_ and rounds integral parameters in {*θ*_***d***_}* to integers while ensuring that |*S*({*θ*_***d***_}*)| ≤ *N*. The probe set pooled across datasets is then *S*({*θ*_***d***_}*).

It is possible that CATCH cannot find a choice of {*θ*_***d***_} such that |*S*({*θ*_***d***_})| ≤ *N*. This might be the case, for example, if the grid of *θ*_***d***_ values over which a user precomputes *s*(*d*, *θ*_***d***_) has too small a range to satisfy the constraint. That is, one or more of the parameter values may need to be relaxed (across one or more datasets) to obtain ≤ *N* probes. When this happens, our implementation of CATCH raises an error and suggests that the user provide less stringent choices of parameter values.

We implemented CATCH in a Python package that is available at https://github.com/broadinstitute/catch.

### Design of viral probe sets presented here

#### Input sequences for design of probe sets

We designed four probe sets using publicly available sequences. The design of V**_ALL_** (356 viral species) incorporated available sequences up to June, 2016; V**_WAFR_** (23 viral species) up to June, 2015; V**_MM_** (measles and mumps viruses) up to March, 2016; and V**_ZC_** (chikungunya and Zika viruses) up to February, 2016. Most sequences we used as input for designing probe sets are genome neighbors (i.e., complete or near-complete genomes) provided in NCBI’s accession list of viral genomes^80^ and were downloaded from NCBI GenBank^36^. We selected a small number of other genomes using the NIAID Virus Pathogen Database and Analysis Resource (ViPR)^81^. Supplementary Table 1 contains links to the exact input (accessions and nucleotide sequences) used as input for each probe set.

In particular, in the input to the design of V**_ALL_** we included all sequences in NCBI’s accession list of viral genomes^80^ for which human was listed as a host, along with all sequences from a selection of additional species (Supplementary Table 1). Since genome neighbors for Influenza A virus, Influenza B virus, and Influenza C virus were not included in the accession list, we included a separate selection of sequences for Influenza A virus that encompass all hemagglutinin and neuraminidase subtypes that infect human (in V**_ALL_**, 8,629 sequences), as well as sequences for Influenza B (376 sequences) and C (7 sequences) viruses. Furthermore, we trimmed long terminal repeats from all sequences of HIV-1 and HIV-2 used as input to both V**_ALL_** and V**_WAFR_**. In V**_ZC_** we included, along with genome neighbors, partial sequences of Zika virus from NCBI GenBank^36^.

#### Exploring the parameter space across taxa

To explore the parameter space in the design of V**_ALL_** and V**_WAFR_**, we varied *m*_***d***_ (number of mismatches) and *e*_***d***_ (cover extension) while fixing all other parameters. We pre-computed probe sets over a grid with *m*_***d***_ in {0, 1, 2, 3, 4, 5, 6} and *e*_***d***_ in {0, 10, 20, 30, 40, 50} when finding optimal parameters. In designing V**_ALL_**, we ran the optimization procedure 1,000 times, each with random starting conditions, and picked the choice of the parameter values from the run with the smallest loss. Supplementary Table 1 lists the selected parameter values of each dataset for each probe set, as well as other fixed parameter values.

#### Design additions for synthesis and probe set data

For synthesis of probes in V**_ALL_**, the manufacturer (Roche) trimmed bases from the 3’ end of probe sequences to fit within synthesis cycle limits. Probe lengths did not change considerably after trimming: of the 349,998 probes in V**_ALL_**, which were designed to be 75 nt, 61% remained 75 nt after trimming and 99% were at least 65 nt after trimming. We did not add PCR adapters for amplification to probe sequences in V**_ALL_**. We did add adapters to probe sequences in V**_WAFR_**, V**_ZC_**, and V**_MM_** (designed to be 100 nt and synthesized with CustomArray); we used two sets of adapters (20 bases on each end), selected by CATCH for each probe to minimize probe overlap as described above. Furthermore, in these three probe sets we included the reverse complement of each designed 140 nt oligonucleotide in the synthesis. The probe sequences of each probe set (with the 20 nt adapters where applicable) are available at https://github.com/broadinstitute/catch/tree/cf500c6/probe-designs.

#### Analysis of probe set scaling with parameter values and input size

In all evaluations of how probe counts grow with respect to an independent variable (Supplementary Fig. 1c, Fig. 1b, and Supplementary Fig. 2), we used genome neighbors from NCBI’s accession list of viral genomes^80^ (downloaded in September, 2017) as input. We trimmed long terminal repeats from HIV-1 sequences. The specific sequences are available at https://github.com/broadinstitute/catch/tree/323b639/hybseldesign/datasets/data. In all of these evaluations, we designed 75 nt probes.

In the plots showing probe counts as a function of parameter values (Supplementary Fig. 1c), we varied only the mismatches (*m*) and cover extension (*e*) parameters using the values shown. We set parameters on the longest common substring (*lcf*) and island of exact match (*i*) to their default values: *lcf* equal to the probe length (75) and *i* = 0. For each pair of parameter values shown, we calculated probe counts across 5 replicates, with the input to each replicate being 300 genomes that were randomly selected with replacement. Shaded regions are 95% pointwise confidence bands.

In the plots showing how probe counts scale with the number of input genomes (Fig. 1b and Supplementary Fig. 2), the “Baseline” approach generates probes by tiling each input genome with a stride of 25 nt and removing exact duplicates. The “Clustering-based” approach generates candidate probes using a stride of 25 nt and deems two probes to be redundant if their longest common substring up to *m* mismatches (shown at *m* = 0 and *m* = 4) is at least 65 nt. It then constructs a graph in which vertices represent candidate probes and edges represent redundancy, and finds a probe set by approximating the smallest dominating set of this graph. For running this clustering-based approach, see the design naively.py executable in our implementation of CATCH. The CATCH approach generates candidate probes using a stride of 25 nt and is shown with parameter values (*m* = 0, *e* = 0), (*m* = 4, *e* = 0), and (*m* = 4, *e* = 50), and all other parameters set to default values. Probe counts for Hepatitis C virus and HIV-1 were calculated and plotted with *n* = {1, 50, 100, 200, 300, …, 1000} input genomes; for Zaire ebolavirus, *n* = {1, 50, 100, 150, …, 850} input genomes; and for Zika virus, *n* = {1, 25, 50, 75, …, 375} input genomes. For each *n*, we calculated probe counts across 5 replicates, with the input to each replicate being *n* genomes that were randomly selected with replacement. Again, shaded regions are 95% pointwise confidence bands.

### Samples and specimens

Human patient samples used in this study (Supplementary Table 2) were obtained from studies that had been evaluated and approved by the relevant Institutional Review Boards (IRBs) or Ethics Committees at Harvard University (Cambridge, Massachusetts), Partners Healthcare (Boston, Massachusetts), Massachusetts Department of Public Health (Jamaica Plain, Massachusetts), Irrua Specialist Teaching Hospital (Irrua, Nigeria), Nigeria Federal Ministry of Health (Abuja, Nigeria), Sierra Leone Ministry of Health and Sanitation (Freetown, Sierra Leone), Nicaraguan Ministry of Health (Managua, Nicaragua), University of California, Berkeley (Berkeley, California), the Ragon Institute (Cambridge, Massachusetts), Hospital General de la Plaza de la Salud (Santo Domingo, Dominican Republic), Universidad Nacional Autónoma de Honduras (Tegucigalpa, Honduras), Oswaldo Cruz Foundation (Rio de Janeiro, Brazil), and Florida Department of Health (Tallahassee, Florida).

Informed consent was obtained from participants enrolled in studies at Irrua Specialist Teaching Hospital, Kenema Government Hospital, the Ragon Institute, Hospital General de la Plaza de la Salud, Universidad Nacional Autónoma de Honduras, Oswaldo Cruz Foundation, and Universidad Industrial de Santander.

IRBs at the Massachusetts Department of Public Health, Florida Department of Health, and Partners Healthcare granted waivers of consent given this research with leftover clinical diagnostic samples involved no more than minimal risk. In addition, some samples from Kenema Government Hospital and Irrua Specialist Teaching Hospital were collected under waivers of consent to facilitate rapid public health response during the Ebola outbreak and also because the research involved no more than minimal risk to the subjects.

The Harvard University and Massachusetts Institute of Technology IRBs, as well as the Office of Research Subject Protection at the Broad Institute of MIT and Harvard, provided approval for sequencing and secondary analysis of samples collected by the aforementioned institutions.

For all clinical and environmental samples, including samples from the 2018 Lassa outbreak, we extracted RNA using the Qiagen QiAmp viral mini kit, except in cases where samples were provided for secondary use as extracted RNA (Supplementary Table 2). Extractions were performed according to manufacturer’s instructions from 140 µL of biological material inactivated in 560 µL of buffer AVL.

Mock co-infection samples were generated by spiking equal volumes of RNA isolated from 2, 4, 6 or 8 viral seed stocks (dengue virus, Ebola virus, Influenza A virus, Lassa virus, Marburg virus, measles virus, Middle East Respiratory Syndrome coronavirus, and Nipah virus) into RNA isolated from the plasma of a healthy human donor, purchased from Research Blood Components. Ebola virus dilution series were generated by adding 1–10^6^ copies of Ebola virus (Makona) to 30 ng or 300 ng of human K562 RNA. All dilutions were prepared and sequenced in duplicate. For samples where the microbial content was uncharacterized — 26 mosquito pools from the United States, human plasma from 25 individuals with acute non-Lassa virus fevers from Nigeria, and human plasma from 25 individuals with suspected Lassa and Ebola virus infections from Sierra Leone — we created sample pools by combining equal volumes of extracted RNA for 5 samples per pool (one mosquito pool contained 6), resulting in 15 final pools (5 mosquito, 5 Nigeria, and 5 Sierra Leone).

### Construction of sequencing libraries

We first removed contaminating DNA by treatment with TURBO DNase (Ambion) and prepared double-stranded cDNA by priming with random hexamers followed by synthesis of the second strand as previously described^14^. We used the Nextera XT kit (Illumina) to prepare sequencing libraries with modifications to enable hybrid capture^10^. Specifically, we used non-biotinylated i5 indexing primers (Integrated DNA Technologies) in place of the manufacturer’s standard i5 PCR primers. As cDNA concentrations from clinical samples are typically lower than the recommended 1 ng, input to Nextera XT was 5 µL of cDNA, except in the case of Ebola serial dilutions where input was 1 ng. Samples underwent 16–18 cycles of PCR and final libraries were quantified using either the 2100 Bioanalyzer dsDNA High Sensitivity assay (Agilent) or by qPCR using the KAPA Universal Complete Kit (Roche). We also prepared sequencing libraries from water with each batch as a negative control.

### Hybrid capture of sequencing libraries

We synthesized the 349,998 probes in V**_ALL_** using the SeqCap EZ Developer platform (Roche). Since the number of features on the array was 2.1 million, we repeated the design 6 times (6 × final probe density). We used these biotinylated single-stranded DNA probes directly for hybrid capture experiments. We performed in solution hybridization and capture according to manufacturer instructions (SeqCapEZ v5.1) with modifications to make the protocol compatible with Nextera XT libraries. Specifically, we pooled up to 6 individual sequencing libraries with at least 1 unique index together at equimolar concentrations (≥ 3 nm) in a final volume of 50 µL. We replaced the manufacturer’s indexed adapter blockers with oligos complementary to Nextera indexed adapters (P7 blocking oligo: 5’-AAT GAT ACG GCG ACC ACC GAG ATC TAC ACN NNN NNN NTC GTC GGC AGC GTC AGA TGT GTA TAA GAG ACA G/3ddC/-3’; P5 blocking oligo: 5’-CAA GCA GAA GAC GGC ATA CGA GAT NNN NNN NNG TCT CGT GGG CTC GGA GAT GTG TAT AAG AGA CAG /3ddC/-3’; Integrated DNA Technologies). The concentration of Nextera XT adapter blockers was reduced to 200 µm to account for sample input < 1 µg. The concentration of probes was also reduced to account for the replication of our V**_ALL_** probe set 6 × across the 2.1 million features. We incubated the hybridization reaction overnight (∼ 16 hrs). After hybridization and capture on streptavidin beads, we amplified library pools using PCR (14–16 cycles) with universal Illumina PCR primers (P7 primer: 5’-CAAGCAGAAGACGGCATACGA-3’; P5 primer: 5’-AATGATACGGCGACCACCGA-3’; Integrated DNA Technologies).

We prepared the focused probe sets (V**_WAFR_**, V**_MM_**, V**_ZC_**) using a traditional probe production approach^82^ in which DNA oligos were synthesized on a 12k or 90k array (CustomArray). To minimize PCR amplification bias and formation of concatemers by overlap extension we performed two separate emulsion PCR reactions (Micellula, Chimerx) to amplify the nonoverlapping probe subsets (assigned adapters *A* and *B* as described above). One primer in each reaction carried a T7 promoter tail (GGATTCTAATACGACTCACTATAGGG) at the 5’ end. We performed *in vitro* transcription (MEGAshortscript, Ambion) on each of these pools to produce biotinylated capture-ready RNA probes. Pools were aliquoted and stored at −80 °C and combined at equal concentration and volume immediately prior to use. Hybrid capture was a modification of a published protocol^82^. Briefly, we mixed the probes, salmon sperm DNA and human Cot-1 DNA, adapter blocking oligonucleotides and libraries and hybridized overnight (∼ 16 hrs), captured on streptavidin beads, washed, and re-amplified by PCR (16– 18 cycles). PCR primers and index blockers were the same as those used in the protocol for the V**_ALL_** probe set. In some cases, we changed the Nextera XT indexes during final PCR amplification to enable sequencing of pre- and post-capture samples on the same run.

We pooled and sequenced all captured libraries on IIlumina MiSeq or HiSeq 2500 platforms. Pre-capture libraries for all samples were also sequenced to allow for comparison of enrichment by capture.

### Depth normalization, assembly, and alignments

We performed demultiplexing and data analysis of all sequencing runs using viral-ngs v1.17.0^83, 84^ with default settings, except where described below. To enable comparisons between pre- and post-capture results, we downsampled all raw reads to 200,000 reads using SAMtools^85^. We performed all analyses on downsampled data sets unless otherwise stated. We chose this number as 90% of all samples sequenced on the MiSeq (among the 30 patient and environmental samples used for validation) were sequenced to a depth of at least 200,000 reads. For those few low coverage samples for which we did not obtain > 200,000 reads, we performed all analyses using all available reads unless otherwise noted (Supplementary Table 3). Downsampling normalizes sequencing depth across runs and allows us to more readily evaluate the effectiveness of capture on genome assembly (i.e., the fraction of the genome we can assemble) than an approach such as comparing viral reads per million. It also allows us to more readily compare unique content (see below). A statistic like unique viral reads per unique million reads can be distorted based on sequencing depth in the presence of a high fraction of viral PCR duplicate reads: sequencing to a lower depth can inflate the value of this statistic compared to sequencing to a higher depth.

We used viral-ngs to assemble genomes of all viruses previously detected in these samples or identified by metagenomic analyses, including the LASV genomes from the 2018 Lassa fever outbreak in Nigeria and the EBOV genomes from the dilution series. For each virus we taxonomically filtered reads against many available sequences for that virus (Supplementary Table 10). We used one representative genome to scaffold the *de novo* assembled contigs (Supplementary Table 3, Supplementary Table 5, Supplementary Table 7). We set the parameters `assembly min length fraction of reference` and `assembly min unambig` to 0.01 for all assemblies. We took the fraction of the genome assembled to be the number of base calls we could make in the assembly divided by the length of the reference genome used for scaffolding. To calculate per-base read depth, we aligned depleted reads from viral-ngs to the same reference genome that we used for scaffolding. We did this alignment with BWA^86^ through the `align and plot coverage` function of viral-ngs with the following parameters: `-m 50000 --excludeDuplicates --aligner options ‘-k 12 -B 2 -O 3’ --minScoreToFilter 60`. We counted the number of aligned reads (unique viral reads) using SAMtools^85^ with `samtools view -F 1024`, and calculated enrichment of unique viral content by comparing number of aligned reads before and after capture. viral-ngs removes PCR duplicate reads with Picard based on alignments, allowing us to measure unique content. We excluded samples where one or more conditions had less than 100,000 raw reads for reasons of comparability. Excluded samples are highlighted in red in Supplementary Table 3.

To assess how the amount of viral content detected increases with sequencing depth (Supplementary Fig. 7b, c), we used data from the Ebola dilution series on 10^3^ and 10^4^ copies. At these input amounts, both technical replicates, with and without capture and in both 30 ng and 300 ng of background, yielded at least 2 million sequencing reads. For each combination of input copies, background amount, technical replicate, and whether capture was used, we downsampled all raw reads to *n* = {1, 10, 100, 1000, 10000, 100000, 200000, 300000, …, 1900000, 2000000 }reads. For each *n*, we performed this downsampling 5 times. We depleted reads with viral-ngs, aligned depleted reads to the EBOV reference genome (Supplementary Table 5), and counted the number aligned, as described above. We plotted the number of aligned reads for each subsampling amount in Supplementary Fig. 7b and c, where shaded regions are 95% pointwise confidence bands calculated across the 5 downsampling replicates.

To analyze the relation between probe-target identity and enrichment (Fig. 3b), we used an Influenza A virus sample of avian subtype H4N4 (IAV-SM5). We assembled a genome of this sample both pre-capture and following capture with V**_ALL_** to verify concordance; we used the V**_ALL_** sequence for further analysis here because it was more complete. We aligned depleted reads to this genome as described above (with BWA using the `align and plot coverage` function of viral-ngs and the following parameters: `-m 50000 –excludeDuplicates--aligner options ‘-k 12 -B 2 -O 3’ --minScoreToFilter 60`). For a window in the genome, we calculated the fold-change in depth to be the fold-change of the mean depth post-capture against the mean depth pre-capture within the window. Here, we used windows of length 150 nt, sliding with a stride of 25 nt. We aligned all probe sequences in V**_ALL_** and V**_WAFR_** designs to this genome using BWA-MEM^86^ with the following options: `-a -M –k 8 -A 1 -B 1 -O 2 -E 1 -L 2 -T 20`; these sensitive parameters should account for most possible hybridizations, and include a low soft-clipping penalty to allow us to model a portion of a probe hybridizing to a target while the remainder hangs off. We counted the number of bases that match between a probe and target sequence using each alignment’s MD tag (this does not count soft-clipped ends), and defined the identity between a probe and target sequence to be this number of matching bases divided by the probe length. We defined the identity between probes and a window of the target genome as follows: we considered all mapped probe sequences that have at least half their alignment within the window, and took the mean of the top 25% of identity values between these probes and the target sequence. In Fig. 3b, we plot a point for each window. We did this separately with probes from the V**_ALL_** and V**_WAFR_** designs.

### Within-sample variant calling

For our comparison of within-sample variant frequencies with and without capture (Fig. 3d, Supplementary Table 6), we used 3 dengue virus samples (DENV-SM1, DENV-SM2, and DENV-SM5). We selected these because of their relatively high depth of coverage, in both pre- and post-capture genomes (Supplementary Table 3); the high depth in precapture genomes was necessary for the comparison. We did not subsample reads prior to this comparison, in order to maximize coverage for detection of rare variants. For each of the three samples, we pooled data from three sequencing replicates of the same pre-capture library prior to downstream analysis. For each of these samples we performed two capture replicates on the same pre-captured library (two replicates with V**_WAFR_** and two with V**_ALL_**), and sequenced, estimated, and plotted frequencies separately on these replicates.

After assembling genomes, we used V-Phaser 2.0, available through viral-ngs^83^,84, to call within-sample variants from mapped reads. We set the minimum number of reads required on each strand (`vphaser min reads each`) to 2 and ignored indels. When counting reads with each allele and estimating variant frequencies, we excluded PCR duplicate reads through viral-ngs. In Fig. 3d, we show frequencies for a variant if it is present at ≥ 1% frequency in any of the replicates (i.e., either the pre-capture pool or any of the replicates from capture with V**_WAFR_** or V**_ALL_**). The plot shows positions combined across the three samples that we analyzed.

We estimated the concordance correlation coefficient (*ρ*_*c*/_) between pre- and post-capture frequencies over points in which each is a pair of pre- and post-capture frequencies of a variant in a replicate. Because we had pooled pre-capture data, each pre-capture frequency for a variant is paired with multiple post-capture frequencies for that variant.

### Metagenomic analyses

We used kraken v0.10.6^87^ in viral-ngs to analyse the metagenomic content of our pre- and post-capture libraries. First, we built a database that included the default kraken “full” database (containing all bacterial and viral whole genomes from RefSeq^88^ as of October 2015). Additionally, we included the whole human genome (hg38), genomes from PlasmoDB^89^, sequences covering selected insect species (Aedes aegypti, Aedes albopictus, Anopheles albimanus, Anopheles gambiae, Anopheles quadrimaculatus, Culex pipiens, Culex quinquefasciatus, Culex tarsalis, Drosophila melanogaster, Varroa destructor) from GenBank^36^, protozoa and fungi whole genomes from RefSeq, SILVA LTP 16 S rRNA sequences^90^, UniVec vector sequences, ERCC spike-in sequences and the human pathogenic viral sequences that were used as input for the V**_ALL_** probe design. The database we created and used is available in three parts. It can be downloaded at https://storage.googleapis.com/sabeti-public/meta dbs/kraken full-and-insects 20170602/[*file*] where [*file*] is: `database.idx.lz4` (642 MB), `database.kdb.lz4` (98 GB), and `taxonomy.tar.lz4` (66 MB).

For mock co-infection samples we ran kraken on all sequenced reads. To confirm that enrichment was successful, we calculated the proportion of all reads that were classified as of viral origin. To compare the relative frequencies of each virus pre- and post-capture with V**_ALL_** and V**_WAFR_**, we calculated the proportion of all viral reads that were classified as each of the 8 viral species. For this we used the cumulative number of reads assigned to each species-level taxon and its child clades, which we term “cumulative species counts”.

For each biological sample, we first subsampled raw reads to 200,000 reads using SAMtools^85^ (except for samples with < 200,000 reads, for which we used all available reads). Then, we removed highly similar (likely PCR duplicate) reads from the unaligned reads with the mvicuna tool through viral-ngs. We ran kraken through viral-ngs and separately ran `kraken-filter` with a threshold of 0.1 for classification. For samples where two independent libraries had been prepared and used for V**_ALL_** and V**_WAFR_**, or where the same pre-capture library had been sequenced more than once, we merged the raw sequence files prior to downsampling. To account for laboratory contaminants we also ran kraken on water controls; we first merged all water controls together, and classified reads as described above. We evaluated the presence and enrichment of viral and other taxa using the cumulative species-level counts, as above. To do so we calculated two measures: abundance, which was calculated by dividing pre-capture read counts for each species by counts in pooled water controls, and enrichment, which was calculated by dividing post-capture read counts for each species by pre-capture read counts in the same sample. For our uncharacterized mosquito pools and human plasma samples from Nigeria and Sierra Leone, after capture with V**_ALL_** we searched for viral species with more than 10 matched reads and a read count greater than 2-fold higher than in the pooled water control after capture with V**_ALL_**. For each virus identified we assembled viral genomes and calculated per-base read depth as described above (Supplementary Fig. 11, Supplementary Table 8). When producing coverage plots, we calculated per-base read depth as described above for known samples, except we removed supplementary alignments before calculating depth to remove artificial chimeras.

### Data availability

Sequences used as input for probe design (Supplementary Table 1) are available in the repository at https://github.com/broadinstitute/catch. Sequences of the probe designs are available at https://github.com/broadinstitute/catch/tree/cf500c6/probe-designs. Viral genomes sequenced as part of this study will be deposited in NCBI GenBank^36^ prior to publication under BioProject accession PRJNA431306 (PRJNA436552 for the 2018 Lassa virus genomes).

**Code availability**

The full source code of CATCH is available at https://github.com/broadinstitute/catch under the terms of the MIT license.

---

**Supplemental Figure 1.**
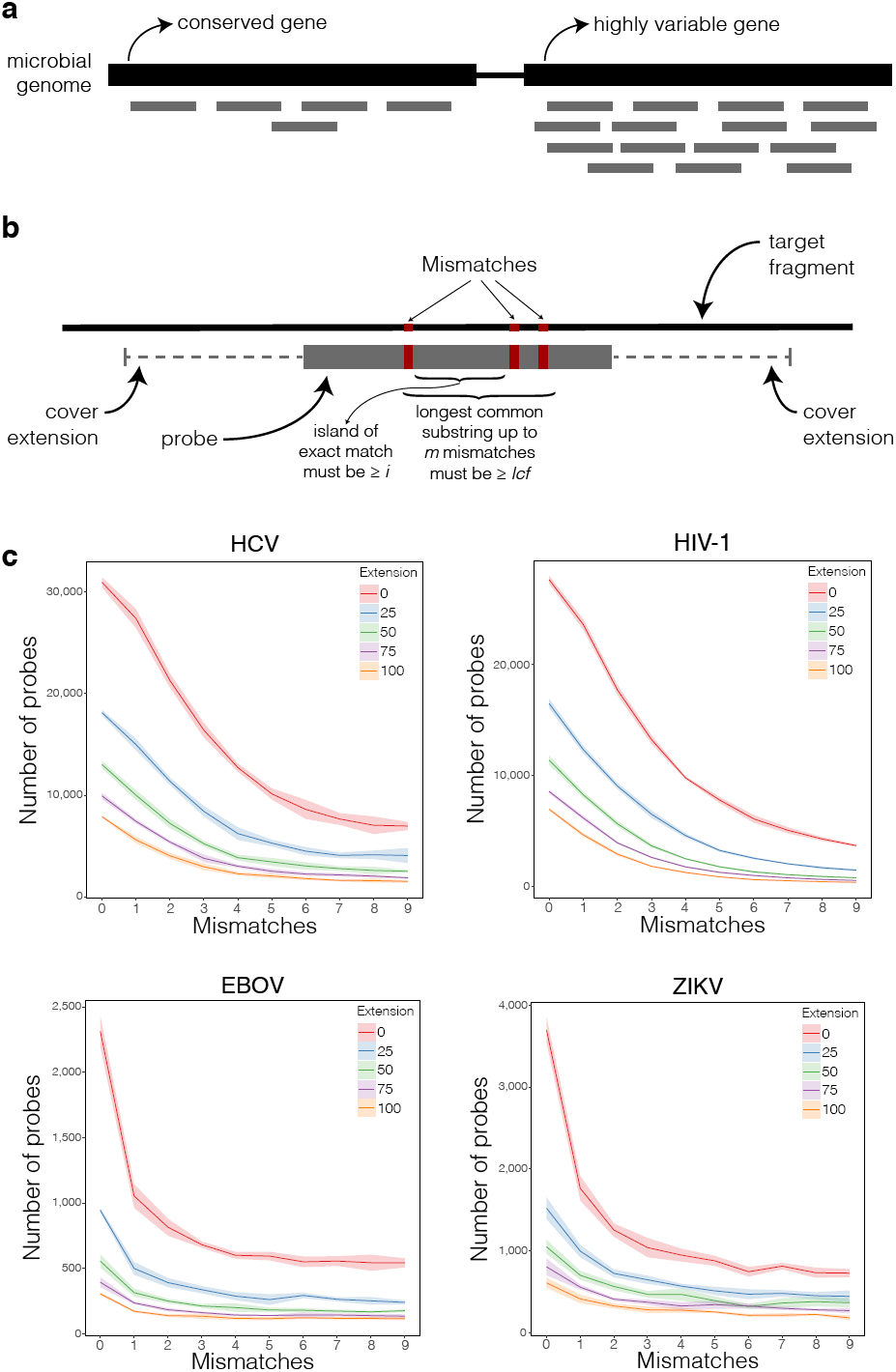
Parameters used by CATCH in default model of hybridization. CATCH models hybridization between each possible candidate probe and the target sequences. Doing so allows CATCH to decide whether a candidate probe captures (or “covers”) a region of the target sequence, and thus find a probe set that achieves a desired coverage of the target sequences under this model. For whole genome enrichment, the desired coverage would typically be 100% of each target sequence. **(a)** Relatively conserved regions (e.g., a particular gene) in the input sequences can be captured with few probes because it is likely that any given probe, under a model of hybridization, will capture observed variation across many or all of the input sequences. Highly variable regions may require many probes to be captured because each given probe may capture the observed variation across only a small fraction of the input sequences. **(b)** By default, CATCH decides whether a probe hybridizes to a region of a target sequence according to the following parameters: a number *m* of mismatches to tolerate and a length *lcf* of a longest common substring. CATCH computes the longest common substring with at most *m* mismatches between the probe and target subsequence, and decides that the probe hybridizes to the target if and only if the length of this is at least *lcf*. If the parameter *i* is provided, CATCH additionally requires that the probe and target subsequence share an exact (0-mismatch) match of length at least *i*. If CATCH decides that the probe hybridizes to the subsequence of the target with which it shares a substring, then it determines that the probe captures the region equal to the length of the probe as well as *e* nt on each side of this region. *e*, termed a cover extension, is a parameter whose value is specified to CATCH, along with *m*, *lcf*, and *i*. Lower values of *m*, higher values of *lcf*, higher values of *i*, and lower values of *e* are more conservative and lead to more probe sequences. (For details, see the description of *f*map in Methods.) **(c)** Number of probes required to fully capture 300 genomes of HCV, HIV-1, EBOV, and ZIKV, for varying values of the mismatches and cover extension parameters, with other parameters fixed. Shaded regions are 95% pointwise confidence bands calculated across randomly sampled input genomes.

**Supplemental Figure 2.**
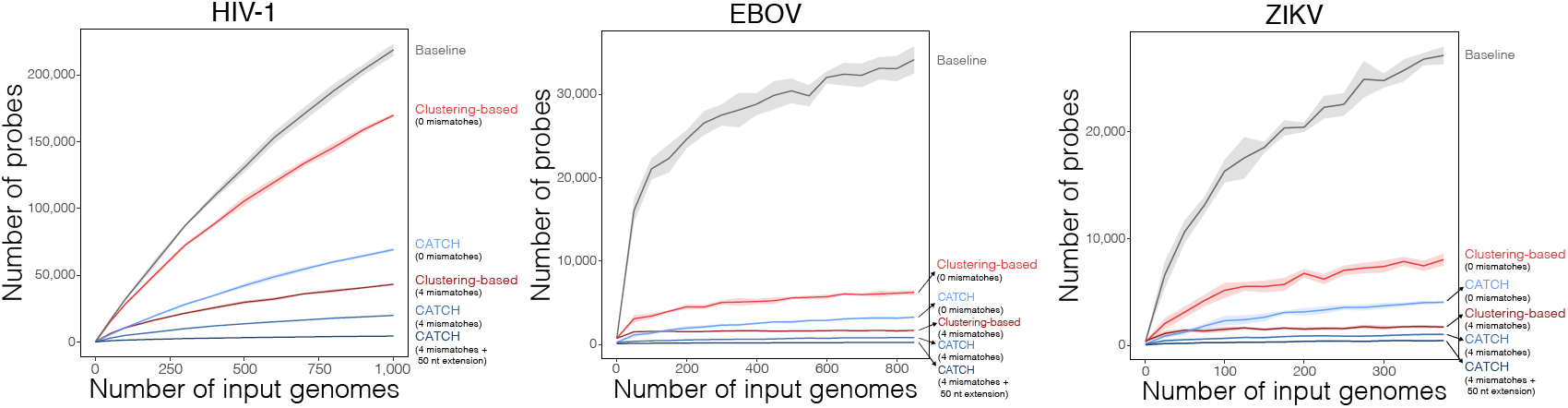
Scaling probe count with diversity of viral genomes. Number of probes required to fully capture increasing numbers of HIV-1, EBOV, and ZIKV genomes. Approaches shown are simple tiling (gray), a clustering-based approach at two levels of stringency (red; see Methods for details), and CATCH at three choices of parameters (blue). Shaded regions are 95% pointwise confidence bands calculated across randomly sampled input genomes.

**Supplemental Figure 3.**
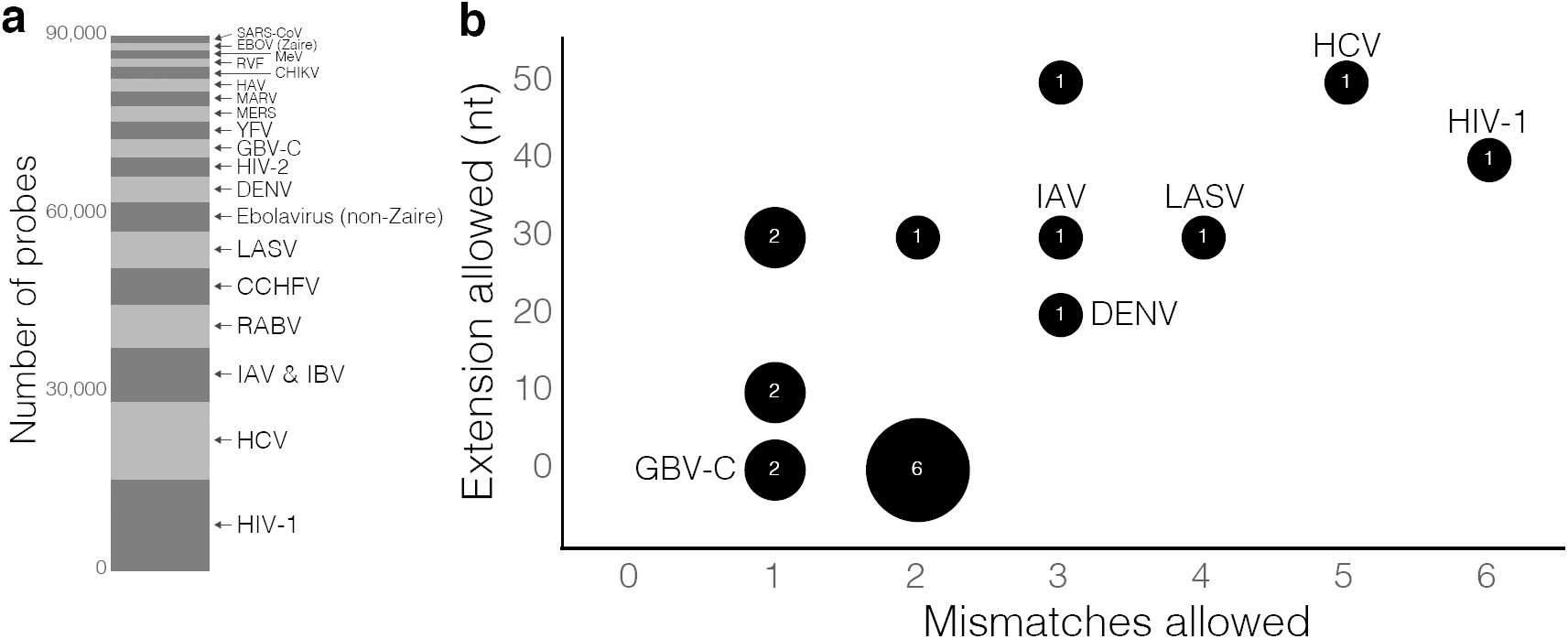
Design of the V**_WAFR_** probe set. **(a)** Number of probes designed by CATCH for each dataset among all 89,990 probes in the V**_WAFR_** probe set. The total includes reverse complement probes, which were added to the design of V**_WAFR_** for synthesis. **(b)** Values of two parameters selected by CATCH for each dataset in the design of V**_WAFR_**: number of mismatches to tolerate in hybridization and length of the target fragment (in nt) on each side of the hybridized region assumed to be captured along with the hybridized region (cover extension). The label within each bubble is the number of datasets that were assigned a particular combination of values. Species included in our sample testing are labeled; for full list of parameter values, see Supplementary Table 1.

**Supplemental Figure 4.**
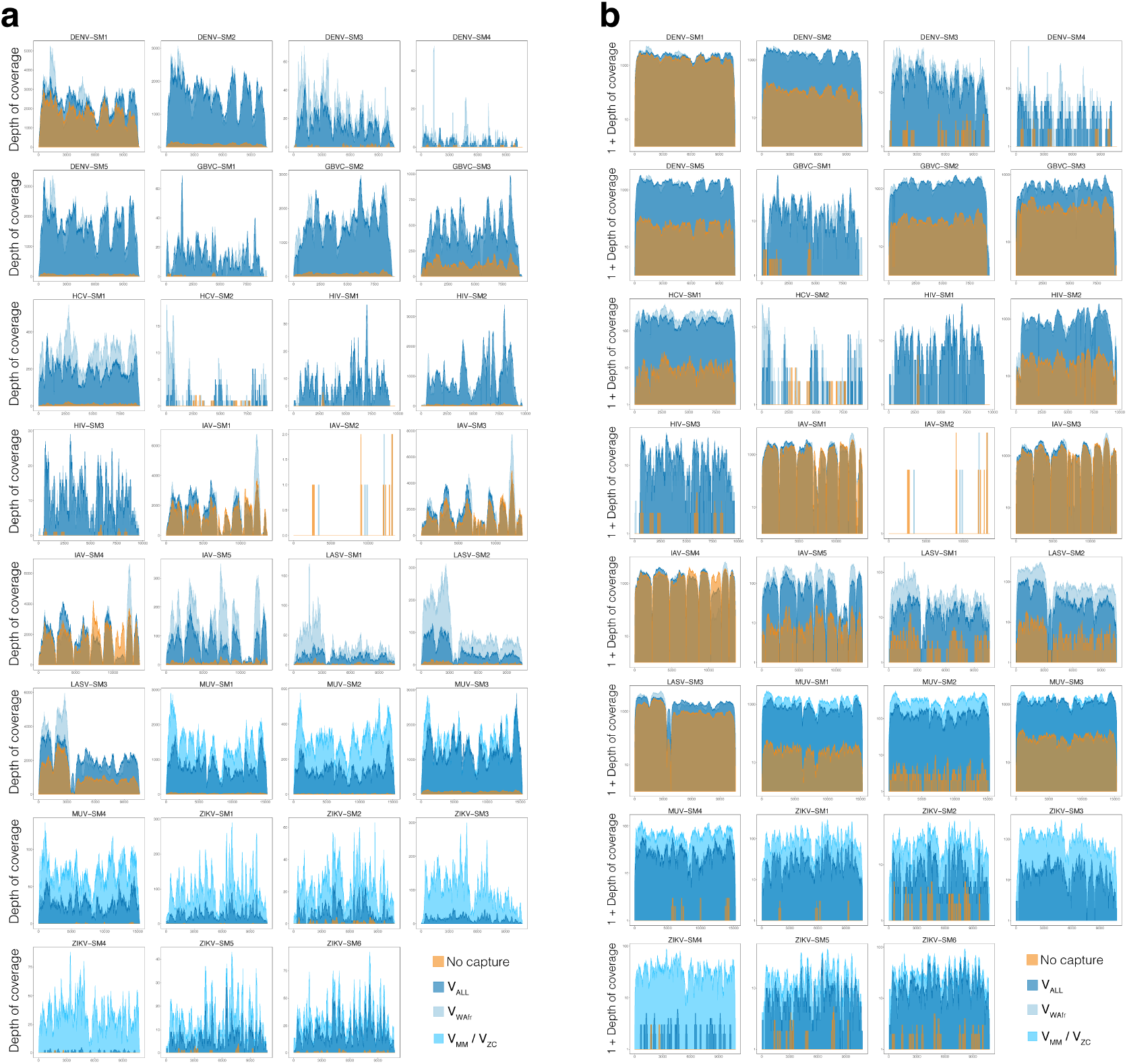
Depth of coverage observed across all viral genomes. Depth of coverage across 31 viral genomes included in this analysis, shown on a **(a)** linear and **(b)** logarithmic scale. The logarithmic scale helps compare variance in depth across each genome between pre- and post-captured data.

**Supplemental Figure 5.**
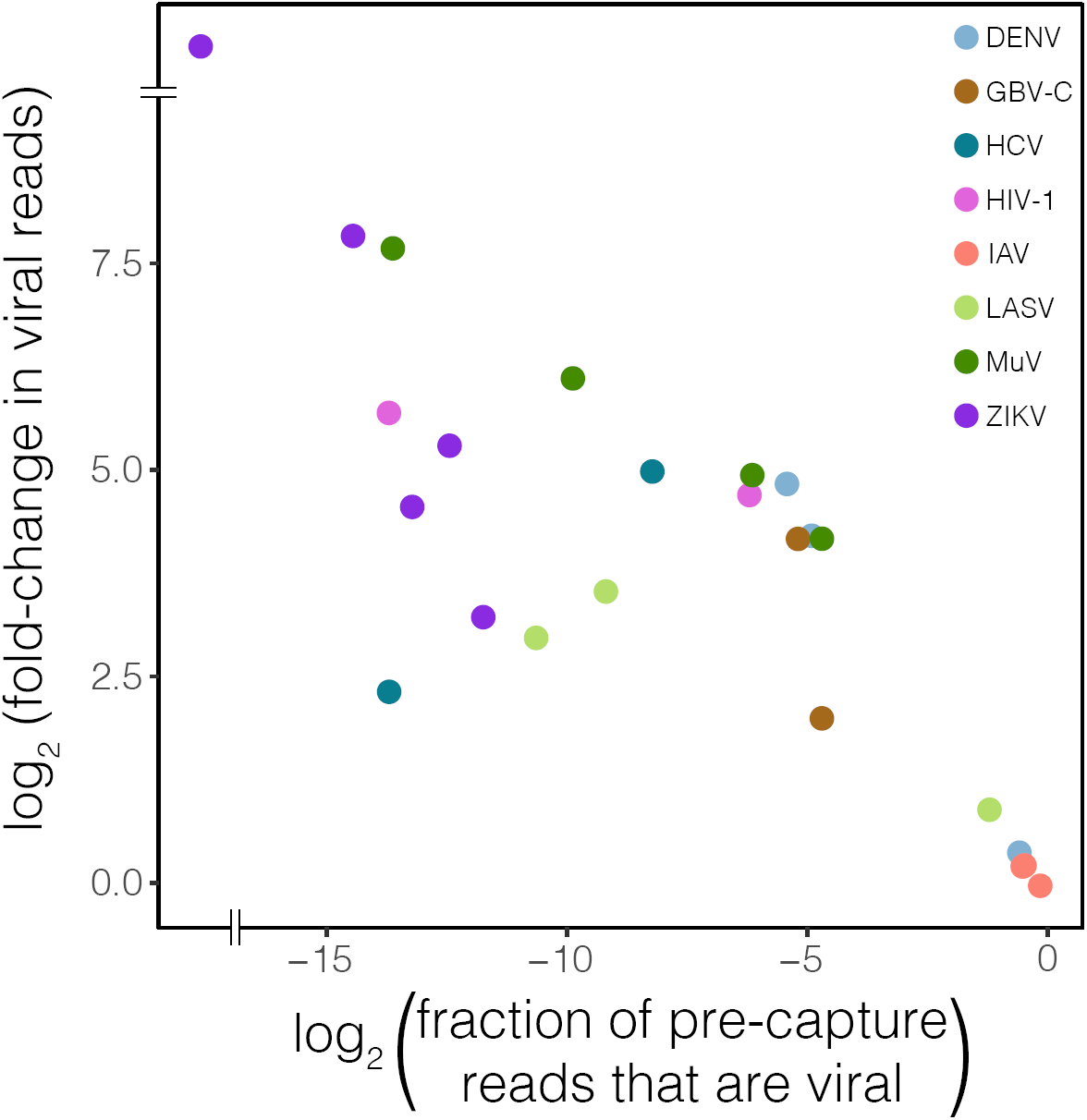
Relation between enrichment of viral content and viral titer. Fraction of all downsampled pre-capture reads that mapped to the reference genome (shown on the horizontal axis) for 24 viral genomes reflects a wide range of initial viral concentrations in these samples. Enrichment (shown on the vertical axis) was calculated by dividing the total number of post-capture reads mapping to a reference genome by the number of mapped pre-capture reads. Those with the highest viral content showed lower enrichment following capture with V**_ALL_**. Seven of the 31 viral genomes included in the analysis are excluded from this plot because they yielded fewer than 200,000 total reads (Supplementary Table 3). Two IAV samples with a high fraction of viral reads pre-capture (bottom right) overlap on the plot. One sample (ZIKV-SM3, top right) showed no viral reads pre-capture, so its fold-change is undefined.

**Supplemental Figure 6.**
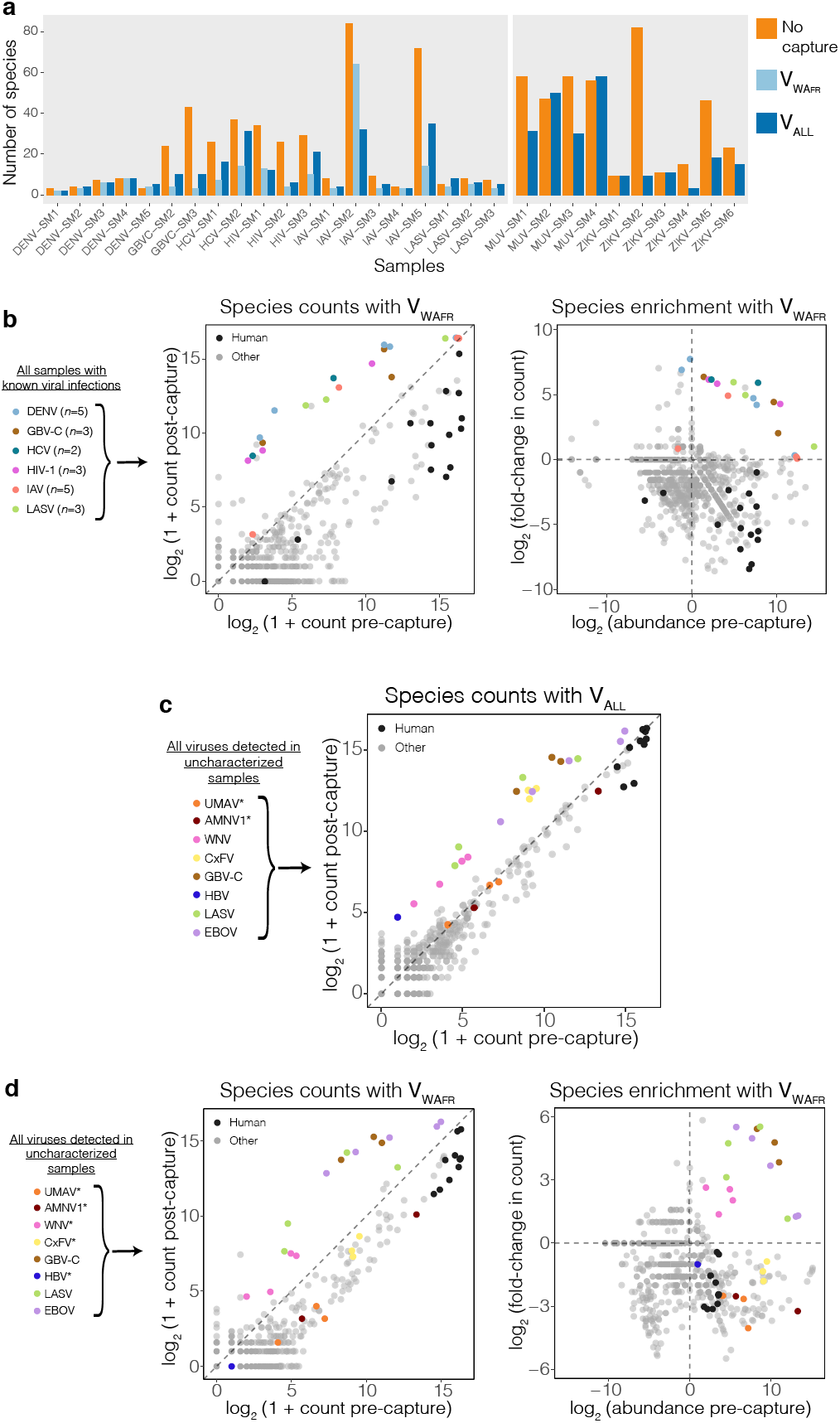
Metagenomic sequencing results for pre- and post-capture samples. **(a)** Number of species detected (with at least 1 assigned read) in samples with known viral infections. Counts are shown before capture (orange), after capture with V**_WAFR_** (light blue), and after capture with V**_ALL_** (dark blue). **(b)** Left: Number of reads detected for each species across samples with known viral infections, before and after capture with V**_WAFR_**. Right: Abundance of each species before capture and fold-change upon capture with V**_WAFR_**. For each sample, the virus known to be present in the sample is colored, and *Homo sapiens* matches in samples from humans are shown in black. **(c)** Number of reads detected for each species across uncharacterized sample pools, before and after capture with V**_ALL_**. Viral species present in each sample (Fig. 4b) are colored, and *Homo sapiens* matches in human plasma samples are shown in black. Asterisks on species indicate ones that are not targeted by V**_ALL_**. **(d)** Same as (b) but for V**_WAFR_** in the uncharacterized sample pools. Asterisks on species indicate ones that are not targeted by V**_WAFR_**. In all panels, abundance was calculated by dividing species counts pre-capture by counts in pooled water controls.

**Supplemental Figure 7.**
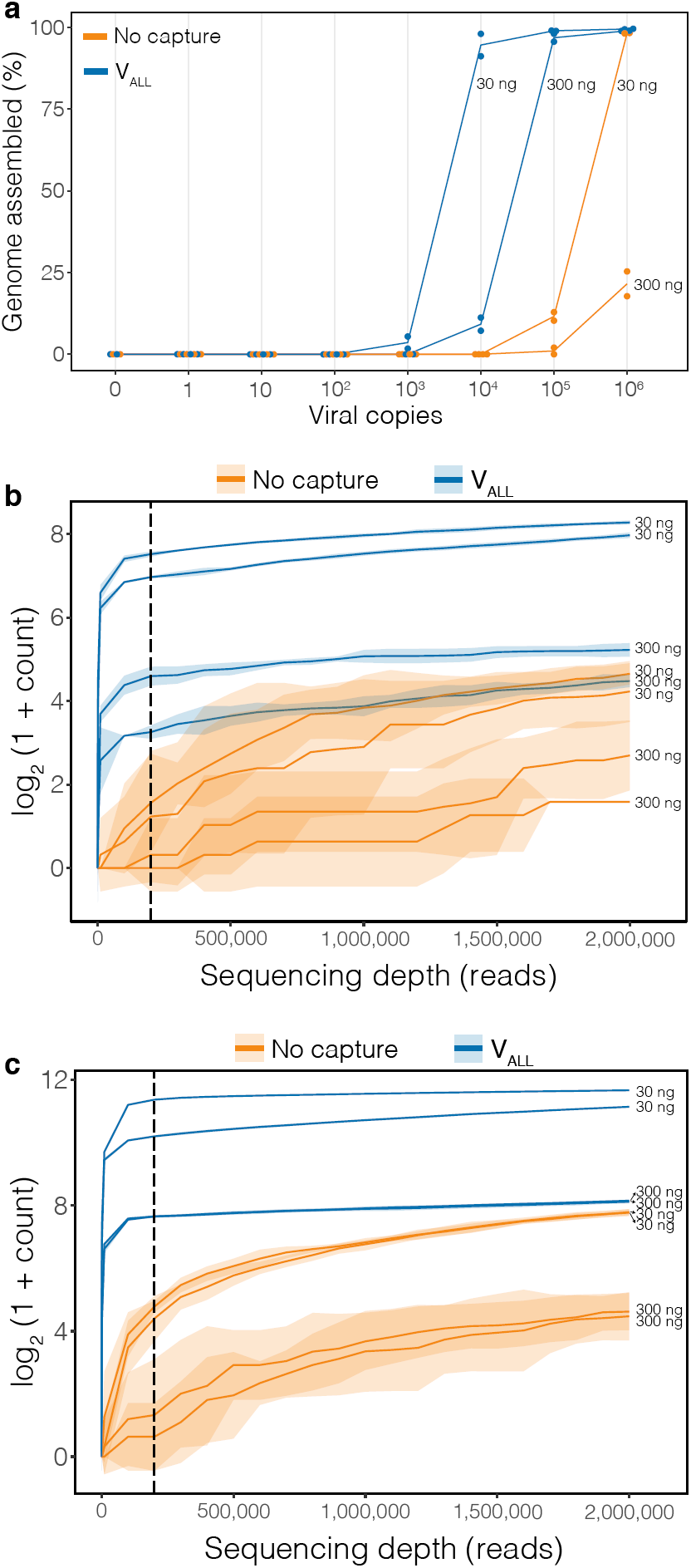
Genome assembly in EBOV dilution series and effect of sequencing depth on amount of viral material sequenced. **(a)** Percent of viral genome assembled in a dilution series of viral input in two amounts of human RNA background. There are *n* = 2 technical replicates for each choice of input copies, background amount, and use of capture (*n* = 1 replicate for the negative control with 0 copies). Each dot indicates percent of genome assembled, from 200,000 reads, in a replicate; line is through the mean of the replicates. Label to the right of each line indicates amount of background material. Assemblies are from read data presented in Fig. 3a. **(b)** Number of unique viral reads sequenced at increasing sequencing depth, from an input of 10^3^ viral copies in different amounts of background. Horizontal axis gives the number of total reads to which a sample was subsampled. Each line is a technical replicate (*n* = 2) and shaded regions are 95% pointwise confidence bands calculated across random subsamplings. Dashed vertical line at 200,000 reads denotes the amount of total reads used in (a) and in Fig. 3a. Viral sequencing data generated after capture with V**_ALL_** saturates more quickly than without capture. **(c)** Same as (b), but from an input of 10^4^ viral copies.

**Supplemental Figure 8.**
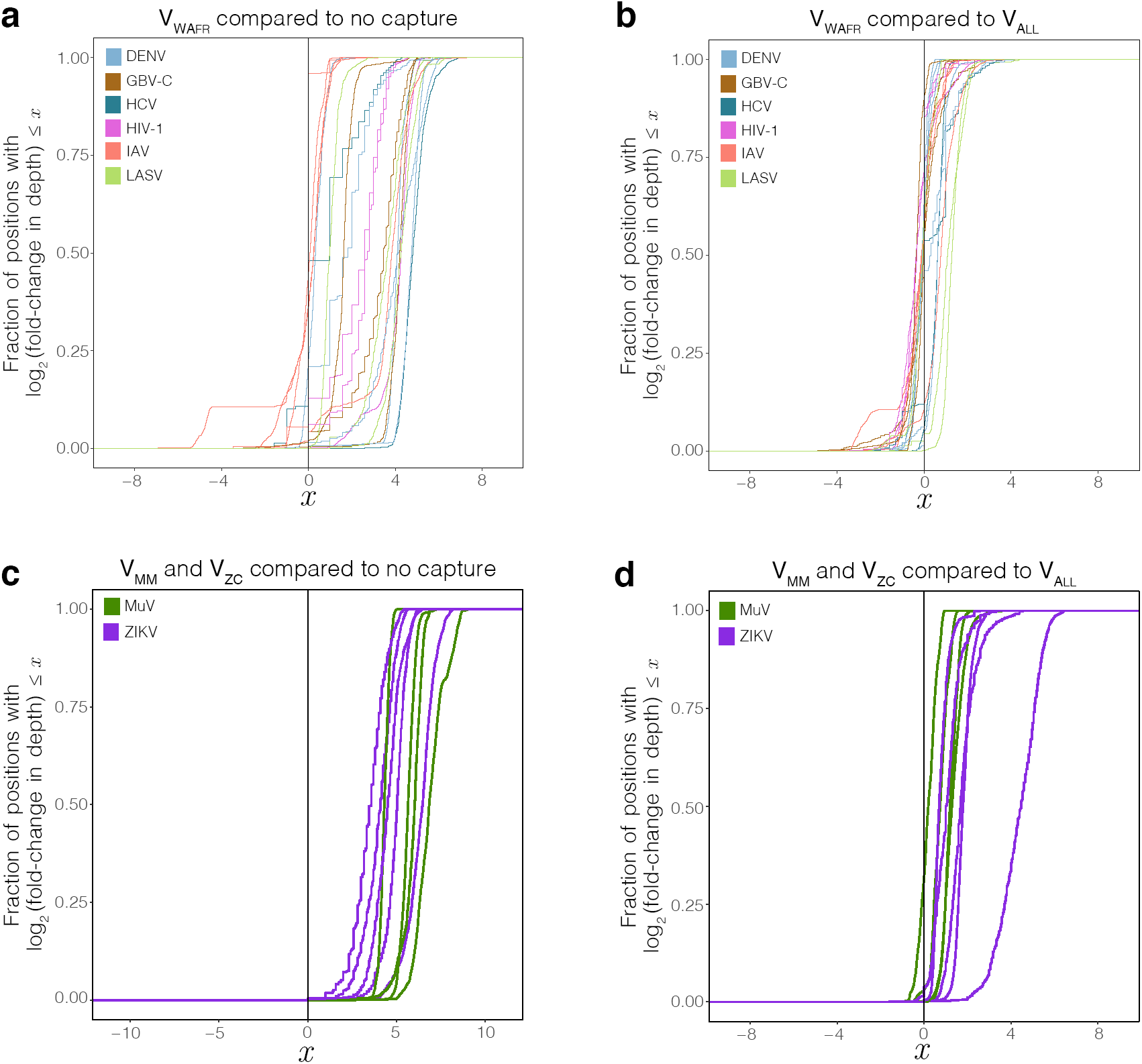
Enrichment in read depth with focused probe sets. **(a)** Distribution of the enrichment in read depth, across viral genomes, provided by capture with V**_WAFR_**. Each curve represents a viral genome. At each position across a genome, the post-capture read depth is divided by the pre-capture depth, and the plotted curve is the empirical cumulative distribution of the log of these fold-change values. **(b)** Distribution of the enrichment in read depth, across viral genomes, provided by V**_WAFR_** over V**_ALL_**. At each position across a genome, the read depth following capture with V**_WAFR_** is divided by the depth following capture with V**_ALL_**, and the plotted curve is the empirical cumulative distribution of the log of these fold-change values. **(c)** Same as (a), but for the two-virus probe sets V**_MM_** and V**_ZC_**. The mumps curves (green) show enrichment provided by V**_MM_** against pre-capture, and the Zika curves (purple) show enrichment provided by V**_ZC_** against pre-capture. **(d)** Same as (b), but for the two-virus probe sets V**_MM_** and V**_ZC_**. The mumps curves (green) show enrichment provided by V**_MM_** against V**_ALL_**, and the Zika curves (purple) show enrichment provided by V**_ZC_** against V**_ALL_**.

**Supplemental Figure 9.**
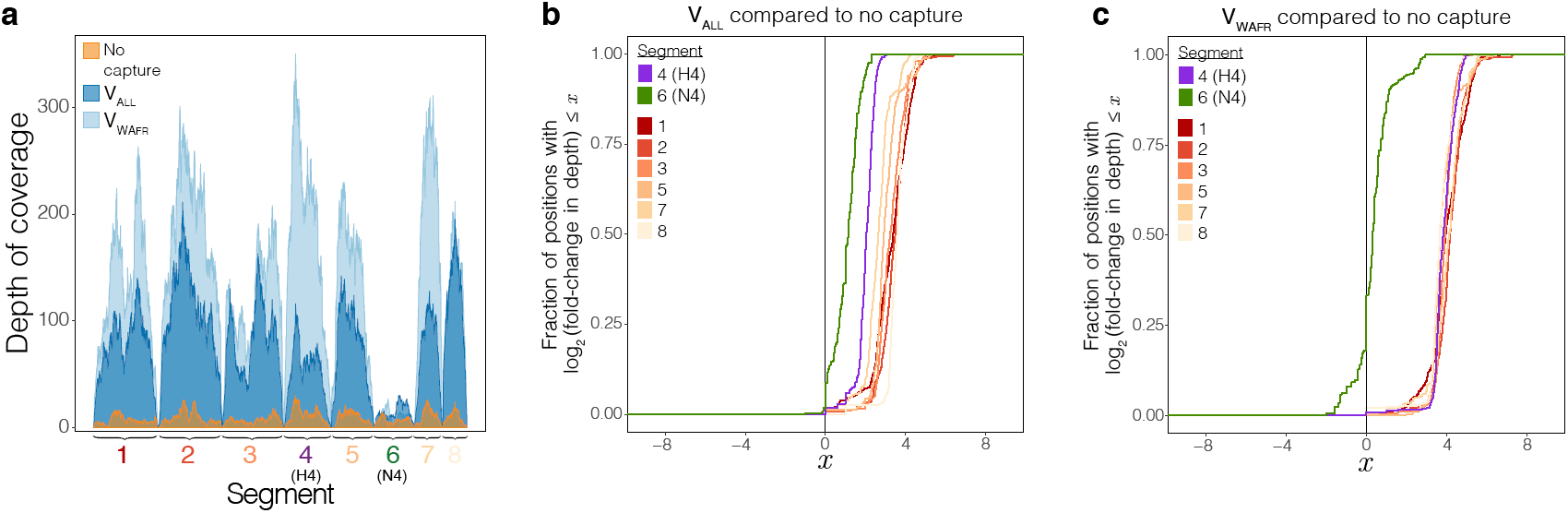
Enrichment across segments of Influenza A virus (H4N4). Variable enrichment across segments of an Influenza A virus sample of subtype H4N4 (IAV-SM5). Segments 4 and 6 contain the most genetic diversity and divergence from probe sequences. No sequences of the N4 subtypes were included in the design of V**_ALL_** or V**_WAFR_**. **(a)** Depth of coverage across the sample’s genome. Each of the eight segments in IAV are labeled. **(b, c)** Distribution of the enrichment in read depth provided by capture with V**_ALL_** (b) and V**_WAFR_** (c). Each curve represents one of the eight segments. At each position across a genome, the post-capture read depth is divided by the pre-capture depth, and the plotted curve is the empirical cumulative distribution of the log of these fold-change values.

**Supplemental Figure 10.**
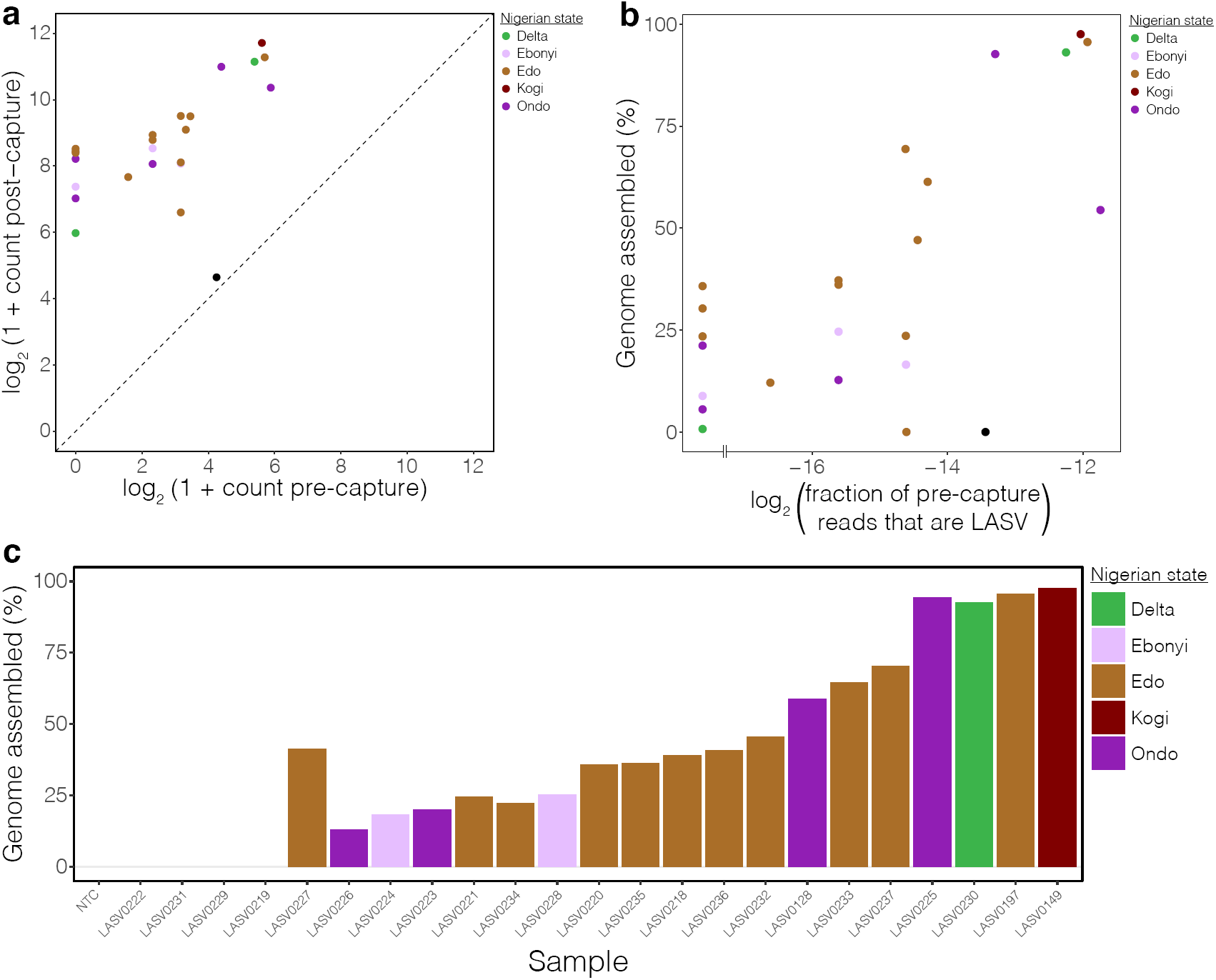
Sequencing results of Lassa virus from the 2018 Lassa fever outbreak in Nigeria. **(a)** Number of unique LASV reads, among 200,000 reads in total, sequenced following capture with V**_ALL_** compared to pre-capture in 23 samples from the 2018 Lassa fever outbreak. Points are colored by the state in Nigeria that the sample is from (black is NTC). **(b)** Percent of LASV genome assembled, after use of V**_ALL_**, against the fraction of pre-capture reads that are LASV. Points to the left of the horizontal break correspond to samples with no LASV reads pre-capture. As in Fig. 4a, reads were downsampled to 200,000 before assembly. Points are colored as in (a). **(c)** Percent of LASV genome assembled, after use of V**_ALL_**. Here, reads were not downsampled before assembly. Bars are ordered as in Fig. 4a and colored by the state in Nigeria that the sample is from.

**Supplemental Figure 11.**
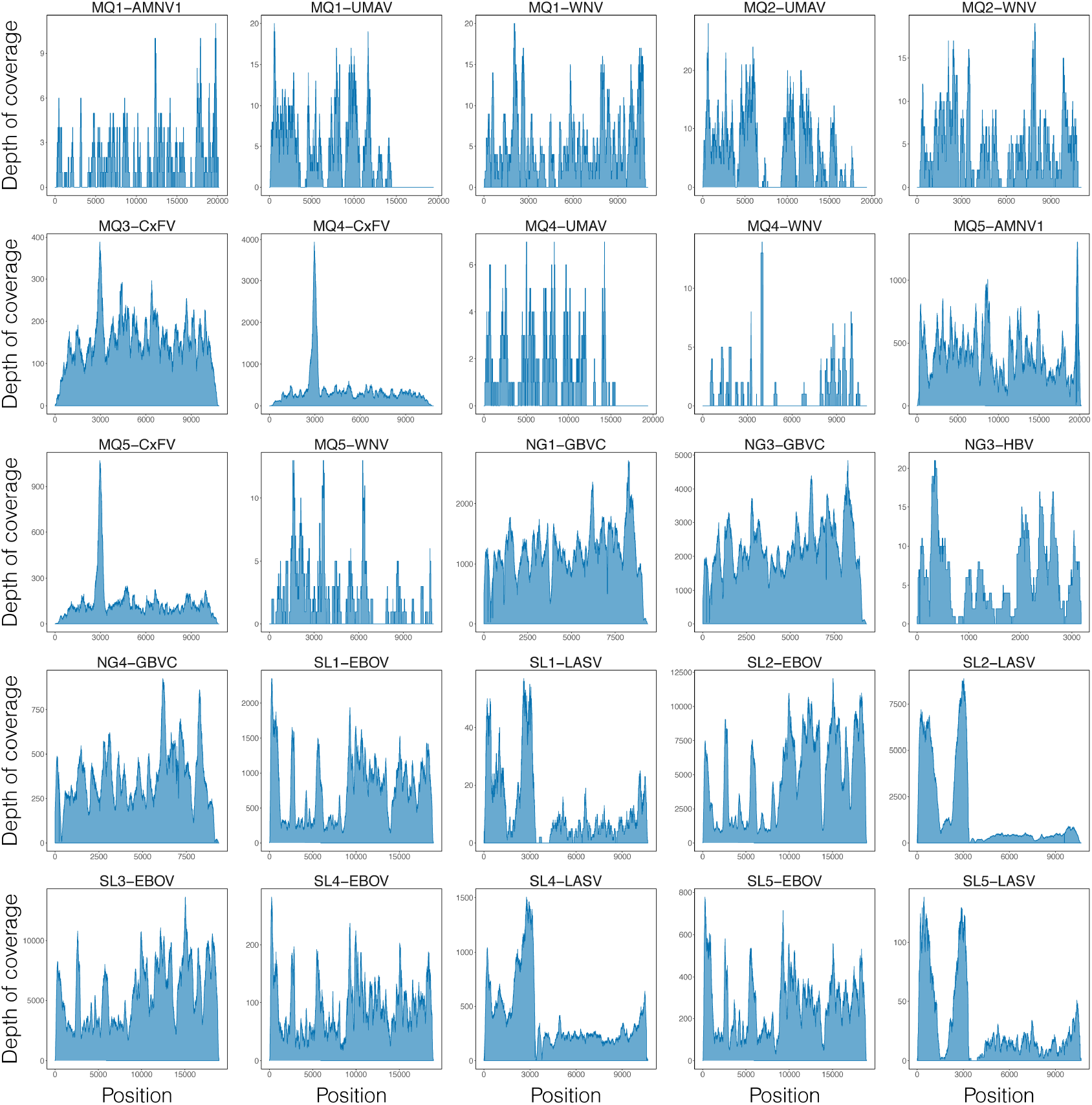
Depth of coverage observed for viral species detected in uncharacterized samples. Depth of coverage plots for 25 viral genomes detected by metagenomic analysis of uncharacterized samples following capture with V**_ALL_** (see Fig. 4b). Read depths are shown on a linear scale.

## List of Supplementary Tables

**Supplementary Table 1**

Input taxa, input data, parameters selected, and other details about the four probe sets presented here.

**Supplementary Table 2**

Origins, source materials, and GenBank accessions for all samples presented here.

**Supplementary Table 3**

Sequencing summary metrics for patient and environmental samples with known viral infections.

**Supplementary Table 4**

Metagenomic species counts for all samples presented here.

**Supplementary Table 5**

Sequencing summary metrics for EBOV dilution series.

**Supplementary Table 6**

Data on within-host variants in DENV samples that were used in the analysis of preservation of within-host variation.

**Supplementary Table 7**

Sequencing summary metrics and metadata for LASV samples from 2018 Lassa fever outbreak in Nigeria.

**Supplementary Table 8**

Sequencing summary metrics for uncharacterized samples.

**Supplementary Table 9**

Cost estimates for sequencing with and without capture.

**Supplementary Table 10**

GenBank accessions used for taxonomic filtering before viral genome assembly.

